# Cellular drivers of injury response and regeneration in the adult zebrafish heart

**DOI:** 10.1101/2021.01.07.425670

**Authors:** Bo Hu, Sara Lelek, Bastiaan Spanjaard, Mariana Guedes Simões, Hananeh Aliee, Ronny Schäfer, Fabian Theis, Daniela Panáková, Jan Philipp Junker

## Abstract

Myocardial infarction is a leading cause of death worldwide, as the adult human heart does not have the ability to regenerate efficiently after insults. In contrast, the adult zebrafish heart has a high capacity for regeneration, and understanding the mechanisms of regenerative processes in fish allows identification of novel therapeutic strategies. While several pro-regenerative factors have been described, the cell types orchestrating heart regeneration remain largely elusive. To overcome this conceptual limitation, we dissected cell type diversity in the regenerating zebrafish heart based on single cell transcriptomics and spatiotemporal analysis. We discovered a dramatic induction of several pro-regenerative cell types with fibroblast characteristics. To understand the cascade of events leading to heart regeneration, we determined the origin of these cell types by high-throughput lineage tracing. We found that pro-regenerative fibroblasts are derived from two separate sources, the epicardium and the endocardium. Mechanistically, we identified Wnt signaling as a key regulator of the endocardial regenerative response. In summary, our results uncover specialized fibroblast cell types as major drivers of heart regeneration, thereby opening up new possibilities to interfere with the regenerative capacity of the vertebrate heart.

---

Heart injury in mammals typically leads to permanent scarring. The zebrafish heart, however, regenerates efficiently after injury, making zebrafish the preeminent model system for heart regeneration (*1, 2*). After cryoinjury, which mimics aspects of myocardial infarction, the injured zebrafish heart undergoes a transient period of fibrosis (*3*), during which the damaged heart muscle regenerates via dedifferentiation and proliferation of cardiomyocytes (*4,5*). Many pathways and factors involved in heart regeneration have been described (6,7,8,9). However, which specific cell types orchestrate the process of heart regeneration remains unclear.

Important molecular signals for zebrafish heart regeneration emanate from the epi- and endocardium (*3,7,10,11*). Other cell types are also required for regeneration: inflammation is an early response to cryoinjury, and depletion of macrophages leads to delayed regeneration (*12,13*). Moreover, fibroblasts are activated upon injury, and ablation of fibroblasts leads to reduced cardiomyocyte proliferation (*14*). All of this suggests that cell types, potentially of a transient nature, residing in the regenerative niche may be important cellular regulators of regeneration. Broadly acting pathways like Wnt tend to influence abundance and expression profiles of multiple cell types in the regenerating heart in potentially complex ways that are difficult to disentangle (*15*). A combination of single-cell analysis and functional experiments is therefore necessary to understand the role of signaling pathways in regeneration.

A number of recent single-cell RNA-sequencing (scRNA-seq) studies have determined cell type diversity in the developing and adult heart (*16*), including large-scale single-cell analysis of the healthy human heart (*17,18*) as well as cell type changes in the mouse after myocardial infarction (*19*). Furthermore, a single-cell analysis of sorted zebrafish cardiomyocytes revealed metabolic changes during the regeneration process (*20*). However, to date no systematic analysis of cardiac cell types in a regeneration-competent organism has been performed. Consequently, our knowledge of the cellular composition of the regenerative niche and the underlying signaling interactions remains incomplete. Hence, current definitions of „ activated” macrophages and fibroblasts rely heavily on transgenes, may be affected by observation bias, and probably underestimate the complexity of transient cell types involved in regeneration. Moreover, the developmental origin of transient cell types, as well as the pathways that generate them, remain unresolved. However, this information is crucial in order to understand the mechanisms underlying generation and activation of pro-regenerative cell types.

## The cellular composition of the regenerating heart

For systematic identification of cardiac cell types in the healthy and regenerating zebrafish heart, we performed scRNA-seq of around 200,000 dissociated cells at different stages pre- and post-injury (Fig. 1A). To limit experimental biases, we did not apply any sorting procedure. In order to include information about the developmental origin of cells, we applied a method for massively parallel lineage tracing based on CRISPR/Cas9 technology (*21,22,23*). By injecting Cas9 and a sgRNA against a multi-copy transgene, we recorded lineage relationships in early development by creating “genetic scars” that serve as lineage barcodes (*21*).

**Figure 1.**
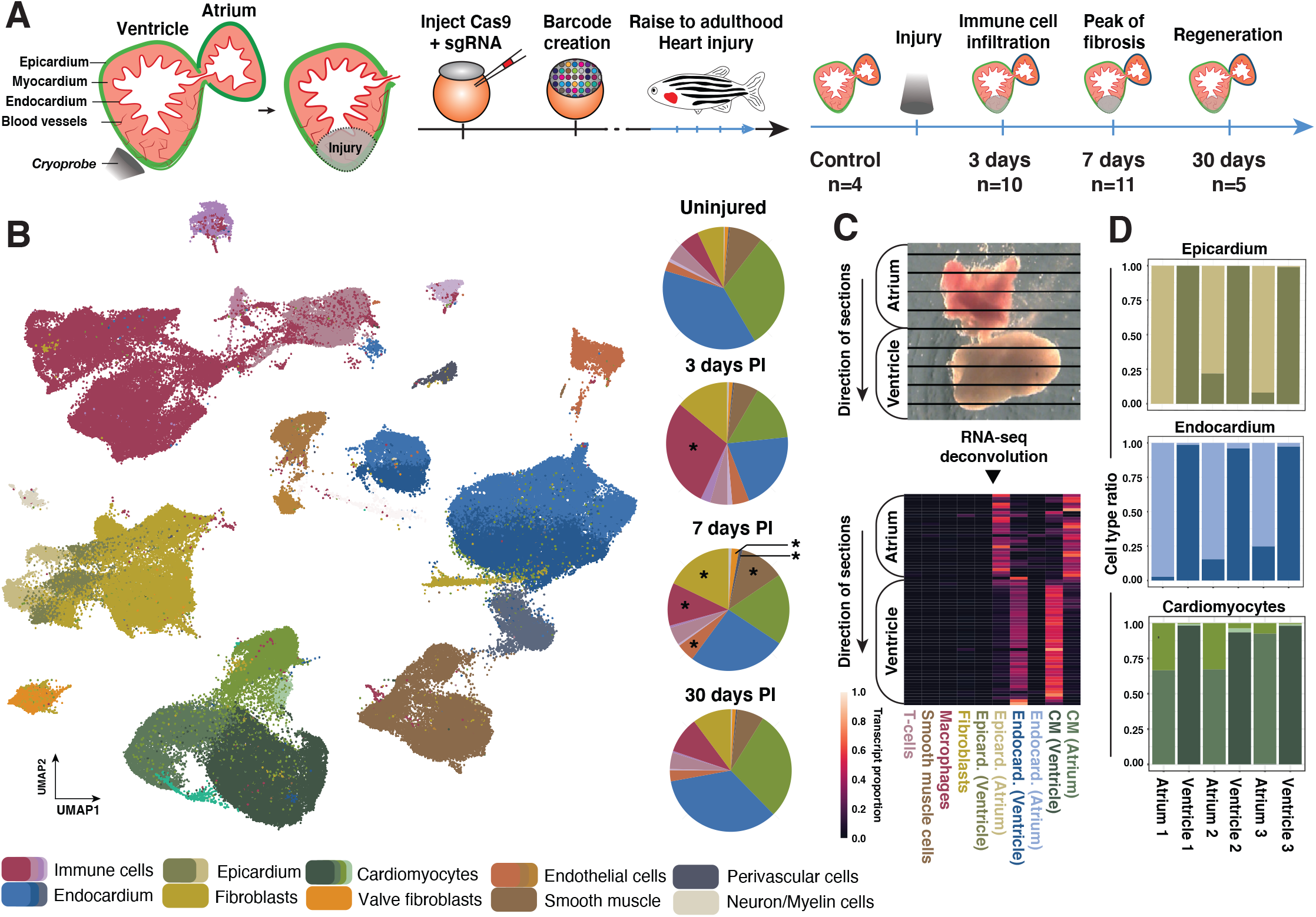
The cellular composition of the regenerating heart. **(A)** Cartoon of the experimental approach. Cells are barcoded during early development, and fish are raised to adulthood. Hearts were harvested either as an uninjured control or at 3, 7 or 30 days post injury (dpi). **(B)** UMAP representation of single-cell RNA-seq data and clustering results. Pie charts show the proportions of different cell types at different time points after injury. In the pie chart representation, similar cell types are grouped and shown by one (representative) color. Asterics (*) denote cell types with a statistically significant change in proportions compared to uninjured control. **(C)** Mapping of single cell data onto a spatially-resolved tomo-seq dataset. A computational deconvolution approach reveals chamber-specific sub-cell-types. **(D)** Distribution of subtypes of cardiomyocytes, endocardial cells, and epicardial cells for scRNA-seq datasets in which atrium and ventricle were physically separated. Color scheme as in Fig. S2.

We first assessed cell type diversity in the healthy and regenerating heart. Clustering of single-cell transcriptomes revealed all major cardiac cell types (Fig. 1B, Fig. S1). As expected, we observed a strong increase in fibroblasts and immune cells after injury (Fig. 1B). Closer inspection of the clustering data revealed a sub-structure among the cell types of the three main layers of the heart – epicardium, myocardium and endocardium (Fig. S2). We hypothesized that this cell type sub-structure might correspond to spatial differences due to functional specification of these cell types in atrium and ventricle. By using the tomo-seq method for spatially-resolved transcriptomics (*24*), and deconvolving this spatial data into single-cell transcriptional profiles (*25*), we could validate atrial and ventricular enrichment for some of these sub-cell-types (Fig. 1C). We confirmed this finding by physical separation of the atrium and ventricle, followed by scRNA-seq (Fig. 1D).

### Identification of cellular drivers of heart regeneration

We identified a further transcriptional sub-structure among the cardiomyocytes (Fig. 2A). In addition to the normal adult cardiomyocytes, which are characterized e.g. by expression of genes involved in ATP synthesis and the tricarboxylic acid cycle (*atp5pd, aldoaa*), we also detected a smaller cluster characterized by genes related to cardiomyocyte development (*ttn, bves, synpo2lb*). These dedifferentiated or newly formed cardiomyocytes, which are a hallmark of the regenerating heart, increased in number already at 3 dpi (days post injury) (Fig. 2A, Fig. S2).

**Figure 2.**
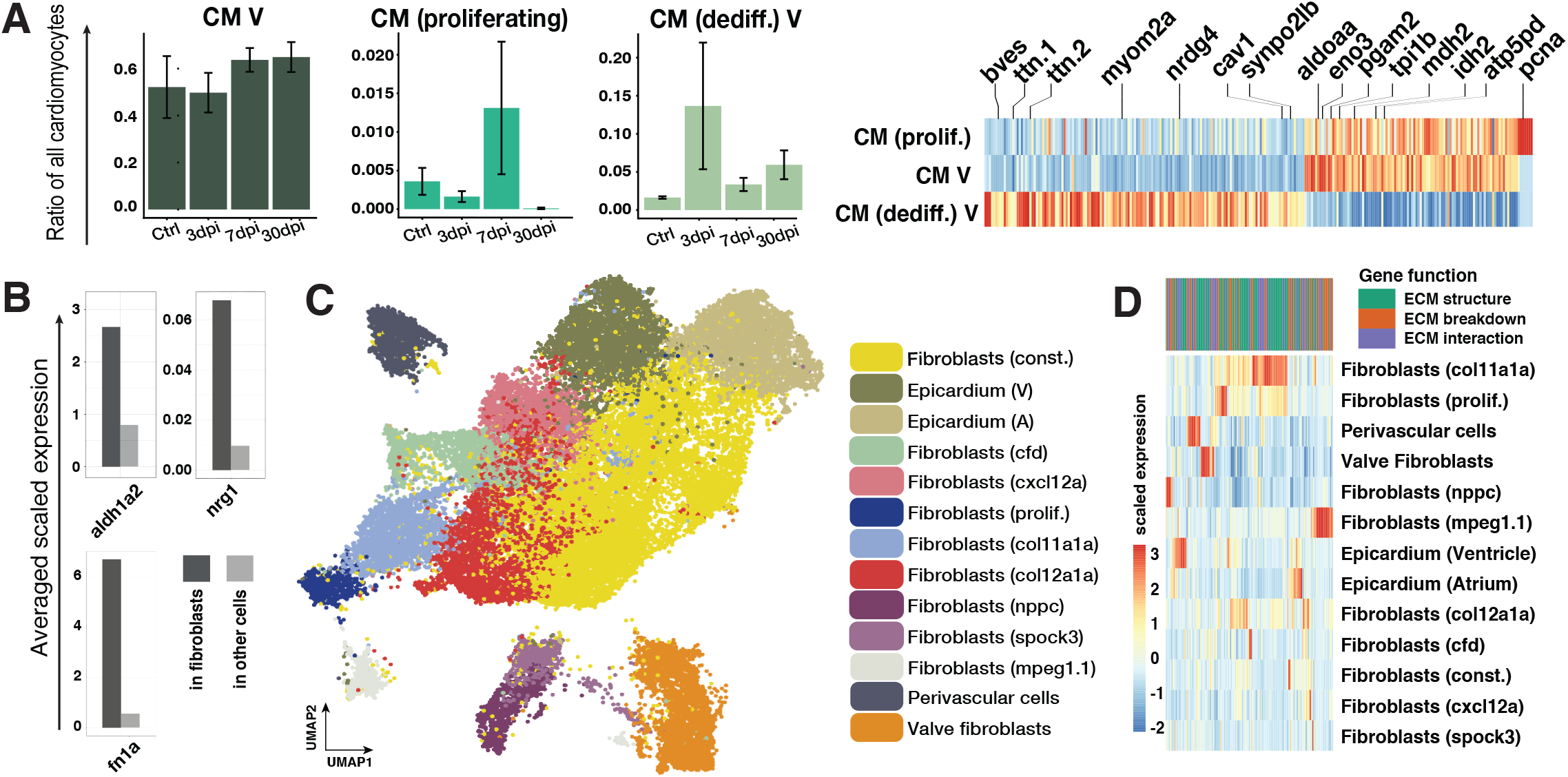
Cell type diversity of cardiac fibroblasts. **(A)** Left: Relative changes of abundance for different subtypes of cardiomyocytes (CM) across the timepoints (error bars show standard error of the mean). Right: Differentially expressed genes between subtypes of cardiomyocytes. **(B)** Comparison of average expression of known pro-regenerative factors in fibroblasts and other cell types. **(C)** UMAP representation ofsubclustering of col1a1a expressing cells. **(D)** Expression of extracellular matrix (ECM) related genes in different fibroblast cell types. The genes are classified according to their contribution to structure, breakdown or interaction of the ECM.

We noticed that three well-established signaling factors in heart regeneration, *aldh1a2* (*6*) (the enzyme producing retinoic acid), the cardiomyocyte mitogen *nrg1* (*26*), and the pro-regenerative extra-cellular matrix (ECM) factor *fn1a* (*27*), are strongly enriched in fibroblasts (Fig. 2B). This prompted us to investigate the diversity of cardiac fibroblasts in more detail. Sub-clustering revealed an unexpectedly large cell type diversity with 13 transcriptionally distinct clusters of fibroblasts (Fig. 2C, Fig. S3), which exhibit pronounced differences in their expression profiles of ECM related genes (Fig. 2D).

To focus our analysis on those fibroblast types that may be part of the regenerative niche, we analyzed the dynamics of these cell types after injury. Three types of fibroblasts, characterized by expression of *col11a1a, col12a1a*, and *nppc*, respectively, stood out as being transiently present at the peak of regeneration (3 and 7 dpi) but virtually absent before injury and after regeneration (Fig. 3A, Fig. S3). Interestingly, *col12a1a*, a non-fibrillar collagen that may act as a matrix-bridging component, has already been shown to be expressed in the epicardial and connective tissues upon heart injury (*28*) and is known to be involved in regeneration of other organ systems in zebrafish (*29*). Two other cell types with an ECM-related function, the perivascular cells as well as the valve fibroblasts, also displayed upregulation after injury. Other fibroblast cell types displayed only moderate changes after injury. Importantly, we noticed that established markers for “activated” fibroblasts like *postnb* (*14*) captured some, but not all fibroblast types that are generated upon injury, and were also expressed in non-fibroblast populations like the epicardial cells (Fig. S4), suggesting that previous marker-based analysis underestimated the transcriptional diversity of the cardiac fibroblast population.

**Figure 3.**
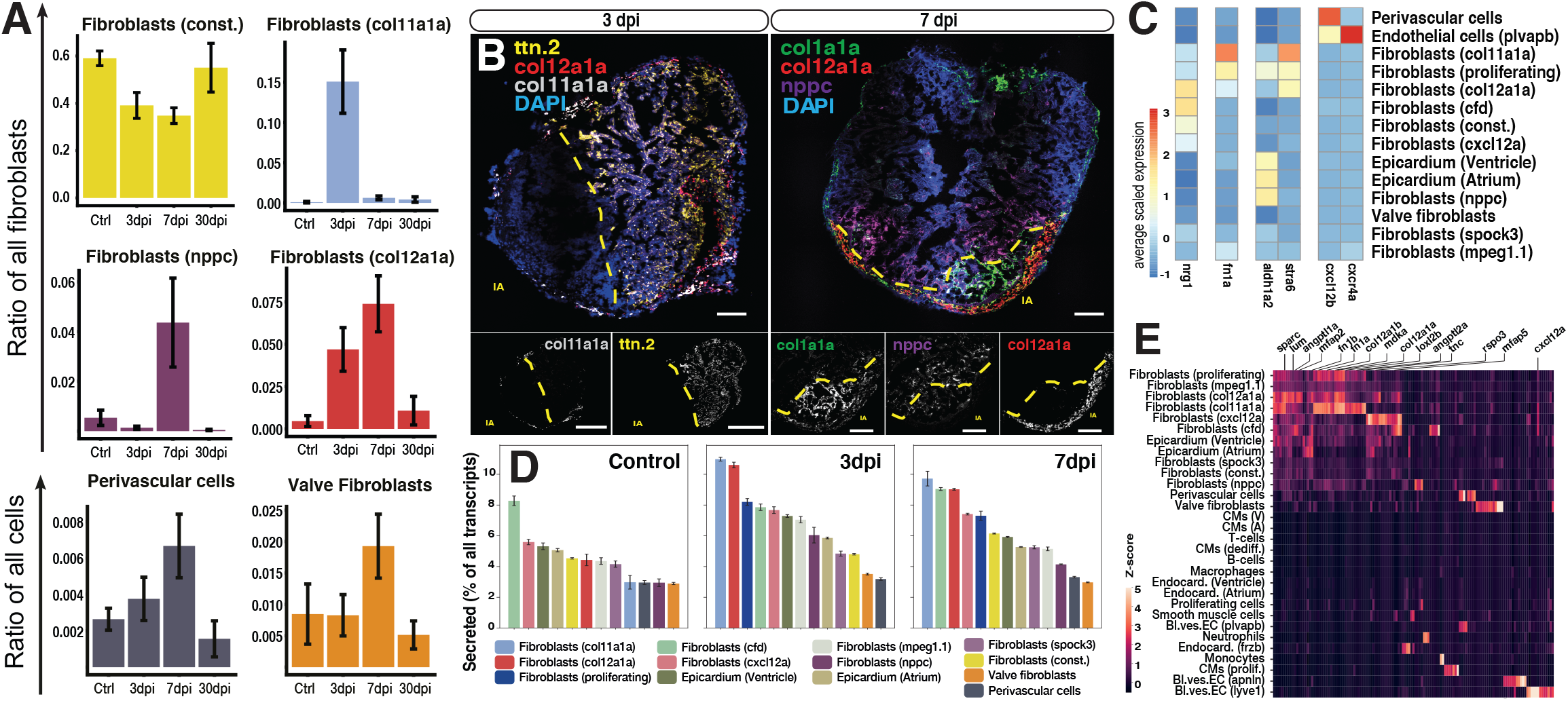
Identification of pro-regenerative cardiac fibroblasts. **(A)** Cell number dynamics of selected fibroblast subclusters, mean value across all replicates of the same timepoint is shown, error bars indicate standard error of the mean. **(B)** Fluorescent in-situ hybridization of marker genes. Left panel: Fibroblasts (const.) (green), Fibroblasts (col12a1a) (red), Fibroblasts (nppc) (purple) at 7 dpi. Right panel: Fibroblasts (col12a1a) (red), Fibroblasts (col11a1a) (white) and dedifferentiated cardiomyocytes expressing ttn.2 (yellow) at 3 dpi. Injury areas are indicated with dotted yellow line. Scale bar: 100*µ*m. **(C)** Average expression of selected signalling genes in fibroblast subclusters. Blood vessel endothelial cells were added to show their interaction with perivascular cells via cxcl12b-cxcr4a signalling. **(D)** Fibroblast cell types, ranked by secretome expression. **(E)** Differentially expressed secretome genes at 3 dpi. Genes with a reported function in regeneration, morphogenesis, tissue development or angiogenesis are highlighted.

To spatially resolve the response of all identified fibroblast cell types, we performed fluorescent *in situ* hybridization (Fig. 3B, Fig. S5, S6). This analysis confirmed the location of the transient fibroblast cell types in the injury area, further corroborating our hypothesis that these cell types contribute to the regenerative niche and regulate injury response and/or regeneration.

We next wanted to understand which of the transient fibroblast cell types express signaling factors involved in heart regeneration (Fig. 3C, Fig. S7). We found particularly high expression of *nrg1* in *col12a1a* fibroblasts, while *fn1a* is expressed almost exclusively in *col11a1a* fibroblasts. Furthermore, we observed that retinoic acid *(aldh1a2)* is produced at high levels by *nppc* fibroblasts as well as epicardium. Interestingly, we observed that the retinoic acid readout gene *stra6* (*30,31*) is expressed highly and specifically in *col11/col12* fibroblasts, but not in cardiomyocytes. Finally, we noticed a very specific interaction between perivascular cells and blood vessel endothelium via *cxcl12b-cxcr4a* chemokine signaling. Perivascular cells are a known regulator of blood-vessel formation (*32*), and it was recently shown that *cxcl12b-cxcr4a* signaling is important for neovascularization of the regenerating heart (*33*).

Finally, we reasoned that the *col11/col12* and *nppc* fibroblasts might express additional secreted factors with a pro-regenerative function in heart regeneration. Indeed, a bioinformatic analysis revealed that expression of secretome genes increases at 3 dpi and 7 dpi compared to uninjured control hearts (Fig. 3D, Fig. S8). Importantly, we found that the *col11/col12* fibroblasts have the highest secretome expression of all detected cell types. The secretome of *col11/col12* fibroblasts is enriched in genes with known functions in regeneration, morphogenesis and tissue development (Fig. 3E, Fig. S8). Interestingly, the secretome at 3 dpi is also enriched in genes with a function in angiogenesis (e.g. *angptl1a* and *angptl2a*), suggesting a role of these cell types in neovascularization. We next performed a ligand-receptor analysis for the identified cell types. We found that the number of putative cell-cell interactions increased drastically after injury and peaked at 3 dpi (Fig. S9), giving rise to a complex network of ingoing and outgoing connections with a noticeable enrichment in fibroblast subtypes. Of note, we detected many incoming interactions for *plvabp* blood vessel endothelial cells, indicating again a potential role of fibroblasts in inducing neovascularization after heart injury..

In summary, based on their gene expression patterns, their timing of appearance, and their spatial position in the injury area, we identified several fibroblast subtypes that appear to be have a pro-regenerative function during heart regeneration: *col11/12* fibroblasts, *nppc* fibroblasts, and perivascular cells.

### The origin of cardiac fibroblasts

We next aimed to elucidate the origin of the transient pro-regenerative fibroblasts in order to better understand their mechanism of activation. To analyze lineage relationships in a high-throughput manner, we used the LINNAEUS method to reconstruct lineage trees for single cells (*21*). In LINNAEUS, cells are marked by heritable DNA barcodes (genetic scars) in reporter genes that are created by Cas9 during early development. The cell-specific accumulation of these scars can then be used to determine lineage relationships between the cells and build a lineage tree. Here, we injected Cas9 into 1-cell stage embryos in order to record lineage relationships during early development, until gastrulation (*21*), that we read out much later, in the adult heart (all lineage trees are shown in Fig. S10-13). By sequencing scars and transcriptome from the same single cells, we can build lineage trees that reveal shared developmental origins of cell types (Fig. 4A). In a lineage tree, all cells in a node share the same developmental ancestor, and any transient cell originating from a cell in a node would be found in the same node. We calculated correlations between cell type ratios in the different tree nodes to determine which cell types are related by lineage (Fig. 4B).

**Figure 4.**
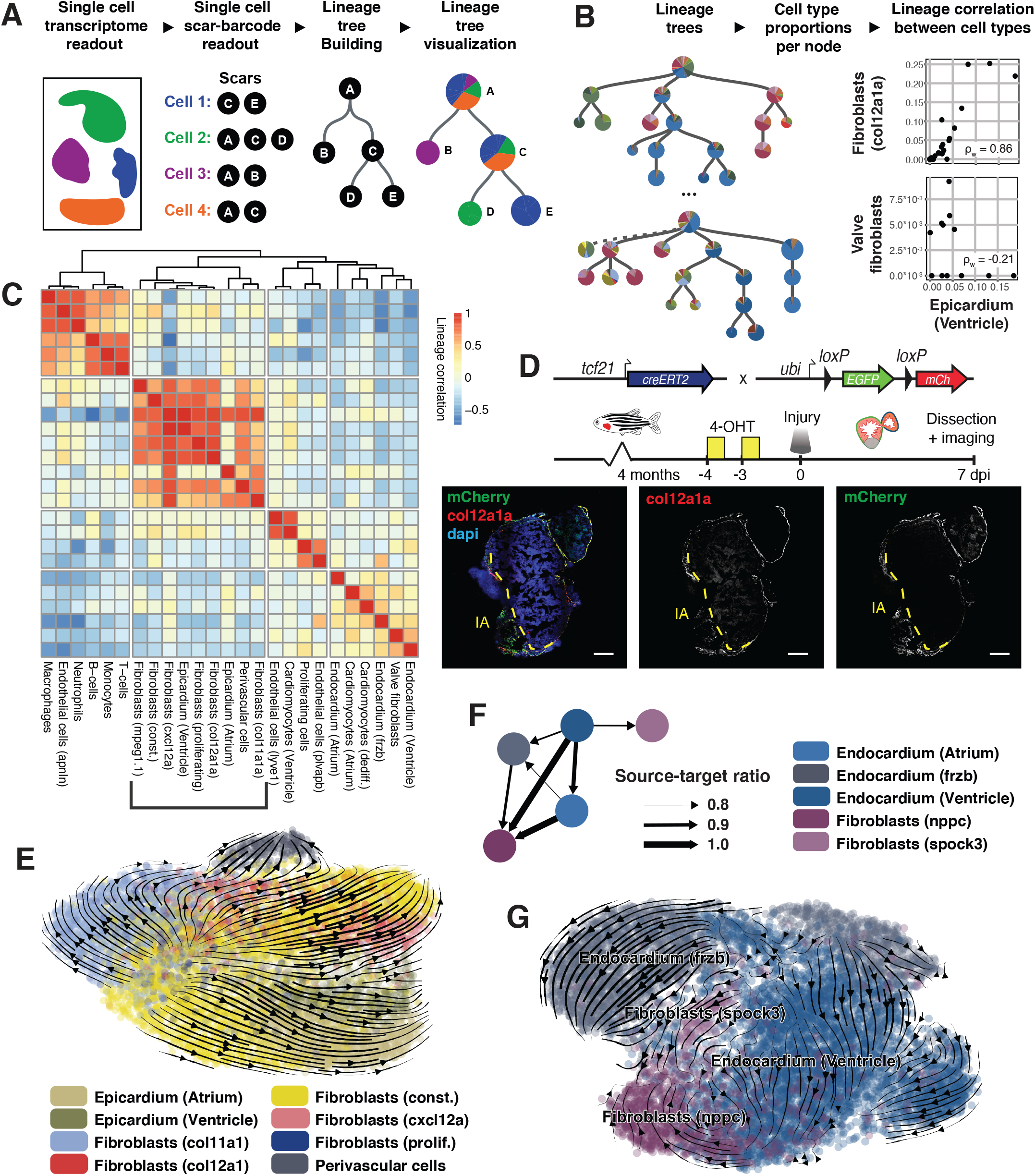
The origin of cardiac fibroblasts. **(A)** Cartoon of lineage tree construction using LINNAEUS. **(B)** We calculate weighted correlations of cell types over the trees in order to quantify lineage similarity. **(C)** Clustering by lineage correlations at 3 dpi reveals epicardial origin of many niche fibroblasts. **(D)** Microsco-py-based lineage tracing confirms epicardial/fibroblast origin of col12a1a fibroblasts. **(E)** Trajectory analysis suggest constitutive fibroblasts as the source of col11a1a and col12a1a fibroblasts at 3 dpi. **(F)** Nppc fibroblasts share asymmetric lineage similarities with endocardial cell types at 7 dpi. **(G)** Trajectory analysis indicates transitions from endocardial cells to spock3 and nppc fibroblasts. Scale bar: 100*µ*m.

Hierarchical clustering of these correlations revealed four clusters of cell types at 3 dpi, the earliest timepoint at which we detect the transient *col11a1a* and *col12a1a* fibroblasts, and seven clusters at 7 dpi, the earliest timepoint at which we detect the transient *nppc* fibroblasts (Fig. 4C, Fig. S14). At both timepoints all immune cells share a common lineage, validating our approach. We observed a clustering of several fibroblast types, including *col11a1a* and *col12a1a* fibroblasts, as well as constitutive fibroblasts, together with the epicardial cells, strongly suggesting that these fibroblast cell types share a developmental origin with the epicardium. Importantly, several fibroblast cell types (*nppc* and *spock3* fibroblasts, valve fibroblasts) were not part of this cluster at 3 dpi and 7 dpi, indicating that these fibroblasts may have a different origin (which we studied in more depth later, in Fig. 4F,G). To validate the lineage origin of *col11a1a* and *col12a1a* fibroblasts, we performed a genetic lineage tracing experiment based on Cre-Lox technology in regenerating hearts using the transgenic line *Tg(tcf21:Cre^ERt2^; ubi:Switch).* Our scRNA-seq data showed that *tcf21* is expressed in epicardial cells, constitutive fibroblasts and other fibroblast types of the epicardial cluster (Fig. S15). After recombination, expression of mCherry colocalizes with expression of *col12a1a* (Fig. 4D), corroborating the origin of the transient *col11a1a* and *col12a1a* fibroblasts from either the epicardium or from epicardial-derived fibroblasts.

LINNAEUS reliably identifies the developmental origin of cell types, but lineage recording is limited to early development. It therefore remains unclear from which source cell type the transient fibroblasts originate upon injury in the adult heart – for instance, we cannot distinguish whether *col11a1a* and *col12a1a* fibroblasts are derived from epicardial cells, constitutive fibroblasts, or any other cell type in the epicardial cluster. Furthermore, expression of *tcf21* is not specific enough to address this question with our Cre-Lox lineage tracing approach (Fig. S15). To further elucidate the origin of the transient fibroblast types, we applied two transcriptome-based trajectory inference methods, partition-based graph abstraction (*34*) (PAGA) and RNA velocity (*35*), to all cells from lineage-related clusters at 3 dpi (Fig. 4E) and 7 dpi (Fig. S16). Of note, when presented with all fibroblast cell types without guidance from lineage trees, PAGA also connects cell types that are not related by lineage according to LINNAEUS, such as *nppc* fibroblasts and epicardial fibroblasts (Fig. S17). This suggests that for processes where several transcriptionally similar cell types are created, integration with explicit lineage tracing methods such as LINNAEUS may be necessary to distinguish real and spurious trajectories. Both timepoints (3 dpi and 7 dpi) show that transient *col11a1a* and *col12a1a*-fibroblasts originate from the constitutive fibroblasts, with an additional contribution from the epicardial cells at 7 dpi (Fig. 4F). Genes that are upregulated along these trajectories are collagens *col11a1a* and *col12a1a*, the pro-regenerative ECM factor *fn1a*, the epicardial activation marker *postnb*, the retinoic acid signaling response gene *stra6,* and the cardiomyocyte mitogen *nrg1* (Fig. S16).

We next focused on the fibroblast types that do not belong to the cluster of epicardium-derived cells, including the transient *nppc* fibroblasts (Fig. 2C). At 7 dpi, we found that the different endocardial cell types (endocardium (Atrium), endocardium (Ventricle) and endocardium (frzb)) were present in over 80% of all nodes containing *nppc* fibroblasts, suggesting a lineage relationship between this fibroblast type and the endocardium (Fig. 4F). Similarly, endocardium (Ventricle) was present in over 80% of nodes with *spock3* fibroblasts. A transcript trajectory analysis revealed transcriptional similarity and a potential differentiation trajectory between *nppc* fibroblasts and the ventricular endocardium (Fig. 4G), confirming their endocardial origin. We observed that *nppc* fibroblasts continue to express endothelial genes (e.g. *vwf, fli1a*) in addition to ECM genes, suggesting that they maintain at least parts of their endocardial gene expression after turning on a fibrotic gene expression program.

Interestingly, the endocardial fibroblasts have low clonality, i.e. only a small fraction of the nodes with endocardial cells also contains endocardial fibroblasts (Fig. S18), suggesting that endocardial fibroblasts are only generated in a subset of the endocardium, probably the injury area. We hypothesized that the internal position of the endocardium, as well as the transient nature and low clonality of the *nppc* fibroblasts, means that these cells are only generated upon an injury that is sufficiently deep to also damage the endocardium. We confirmed that longer contact of the cryoprobe and the heart led to deeper injuries. Indeed, longer exposure to the cryoprobe resulted in much stronger *nppc* expression beyond the border zone, as opposed to injuries with shorter contact time (Fig. S18).

In summary, we combined massively parallel lineage tracing and trajectory inference of single-cell transcriptomes in order to systematically identify the origin of cell types in the regenerative niche. Our analyses revealed a clear separation of epicardial and endocardial-derived fibroblasts, with both lineages giving rise to transient fibroblasts upon injury. While the epicardial origin of cardiac fibroblasts has been clearly established in zebrafish, the existence of atrial and ventricular fibroblasts of endocardial origin had previously not been fully resolved (*11,14*). Given the endocardial origin of these cells, their expression of *aldh1a2* (Fig. 3B), and their localization in the injury border zone (Fig. 3C), we conclude that the *nppc* fibroblasts contribute significantly to retinoic acid signaling in the regenerative niche, which is in line with the previously described endocardial contribution to retinoic acid signaling^6^.

### Cellular dissection of the role of canonical Wnt signaling

While we established the pro-regenerative role of *col11, col12* and *nppc* fibroblasts based on gene expression data, and identified their origin based on lineage analysis, it remains unclear which signaling pathways are required to generate these transient fibroblast cell types upon injury. Functional pathway inhibition experiments are required to determine mechanisms of fibroblast activation. We noticed that fibroblasts express many genes related to Wnt signaling (ligands, receptors, modulators) (Fig. 5A, Fig. S19), which inspired us to investigate the role of canonical Wnt signaling in this system. The role of Wnt signaling in heart regeneration remains an important open question (*15*): On the one hand, Wnt is generally considered a pro-proliferative factor, and Wnt activation has been shown to be beneficial for zebrafish fin and spinal cord regeneration (*29*),(*36*). On the other hand, it was recently reported that activation of Wnt signaling suppressed cardiomyocyte proliferation after ventricular apex resection (*37*). With our knowledge of cardiac cell types as well as their origin, location and dynamics upon injury, we sought to dissect the potentially complex effects of Wnt on the cellular level.

**Figure 5.**
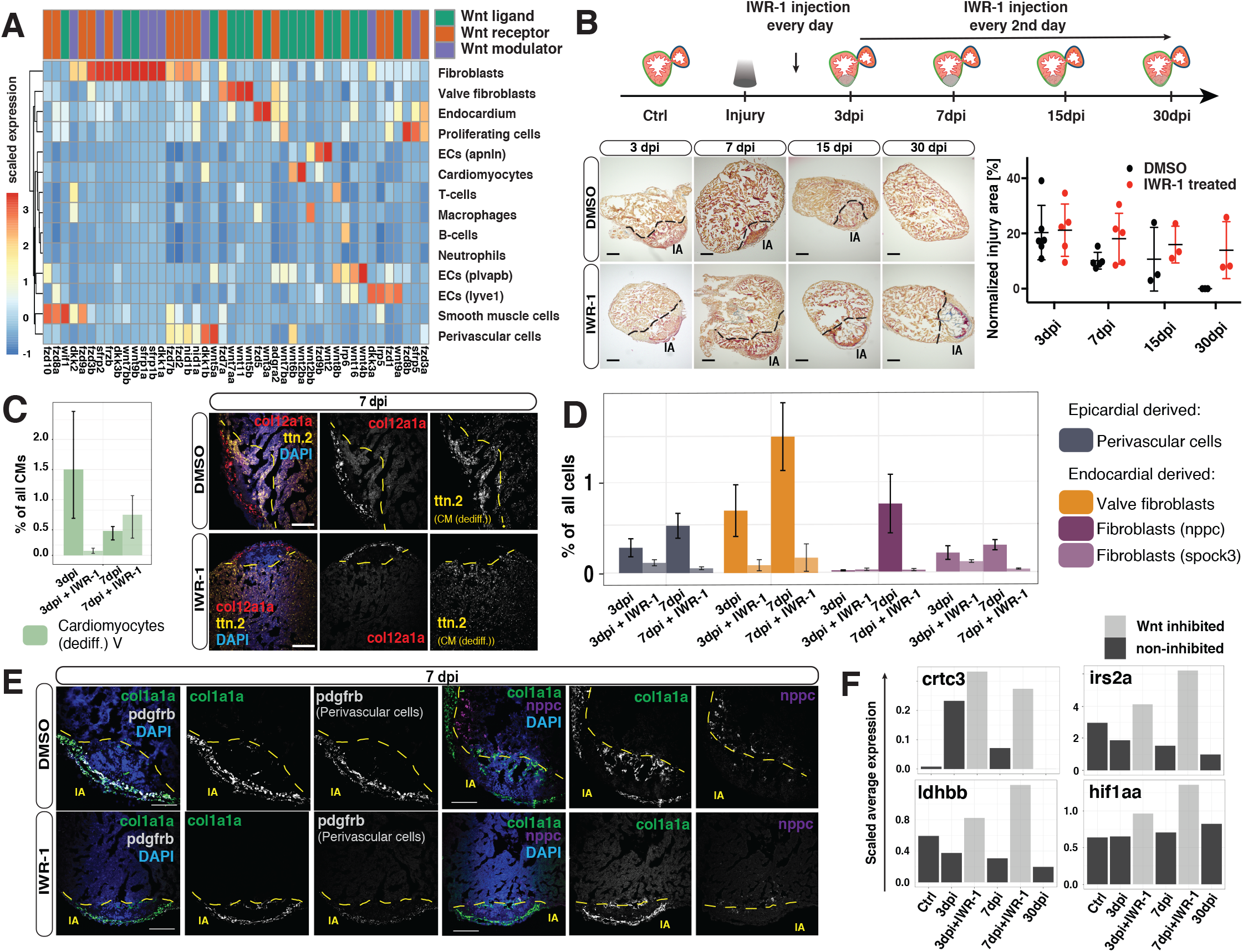
Cellular dissection of the role of canonical Wnt signaling. **(A)** Expression of Wnt signaling factors in the different cell types of the zebrafish heart. **(B)** Upper panel: Cartoon summary of IWR-1 Wnt inhibition experiments. Lower left: Histological comparison of the injury area (IA) at different time points after injury with or without Wnt inhibition. Scale bar: 300*µ*m. Lower right: Relative size of the injury area across all histological replicates, mean and standard deviation are shown. **(C)** Left: Changes in number of dedifferentiated cardiomyocytes at 3 and 7 dpi between Wnt inhibited and control hearts. Right: Localization of dedifferentiated (ttn.2) cardiomyocytes at 7 dpi in the injured heart with or without Wnt inhibiton. **(D)** Changes in abundance of non-cardiomyocte cell types upon Wnt inhibition at 3 and 7 dpi. (in (C) and (D), error bars show standard error of the mean) **(E)** Fluorescent in-situ hybridization of perivascular cells (white) and fibroblasts (nppc) (purple) in injured hearts at 7 dpi with or without Wnt inhibition. (in (C) and (E), scale bar: 100*µ*m) **(F)** Expression of hypoxia induced genes at 3 and 7 dpi with and without Wnt inhibition.

We therefore inhibited canonical Wnt signaling after cryoinjury by using the well-characterized Wnt/β-catenin-dependent signaling inhibitor IWR-1 and observing its effects at 3, 7, 15, and 30 dpi (Fig. 5B). Wnt/β-catenin signaling inhibition led to a pronounced delay in heart regeneration, with prolonged fibrosis and increased injury area compared to the control (Fig. 5B), as previously reported (*37*). Single-cell transcriptomics of IWR-1-treated hearts at 3 dpi and 7 dpi revealed that cardiomyocyte dedifferentiation was delayed (Fig. 5C). Compared to control samples, dedifferentiated cardiomyocytes were reduced at 3 dpi and did not localize to the injury area at 7 dpi (Fig. 5C).

Intriguingly, the perivascular cells, as well as all endocardium-derived fibroblast types (*nppc* fibroblasts, *spock3* fibroblasts, valve fibroblasts) were strongly reduced upon Wnt inhibition, while all other fibroblast cell types were largely unaffected (Fig. 5D, Fig. S20). We confirmed this finding by fluorescent *in situ* hybridization (Fig. 5E). This observation strongly suggests that Wnt signaling is required for the activation of endocardial fibroblasts, akin to the described role of Wnt in inducing endothelial-to-mesenchymal transition in mice (*38,39*). We next examined the consequences of the depletion of perivascular cells. Based on high expression of the chemokine *cxcl12b* in perivascular cells (Fig. 3B), we hypothesized this cell type to be a major driver of neovascularization. Indeed, we observed that hypoxia markers are upregulated in IWR-1 treated hearts upon injury (Fig. 5F), which may indicate a tissue response to the loss of perivascular cells. In summary, we used inhibition of Wnt/β-catenin signaling as a paradigm to dissect the function of complex signaling pathways on the single-cell level. We discovered that Wnt/β-catenin signaling is essential for the transient upregulation of endocardial-derived fibroblasts and perivascular cells, the consequences of which are in agreement with the observed defects in cardiomyocytes response and impaired regeneration.

## Discussion

Here we used a sorting-free approach to determine the cellular composition of the regenerative niche in the zebrafish heart based on single-cell transcriptomics as well as spatio-temporal analysis. This allowed us to identify three transient cell types with fibroblast characteristics (*col11/12* fibroblasts, *nppc* fibroblasts) that are major sources of known pro-regenerative genes like *nrg1, fn1a* and retinoic acid and also express additional secreted factors with a potential pro-regenerative function. These fibroblast cell types are ideally suited as cellular drivers for regeneration based on their expression of pro-regenerative factors, their location in the injury area, and their transient appearance after injury. By combining high-throughput lineage tracing with RNA velocity and single-cell trajectory inference, we were able to systematically identify the origin of these cell types. This approach effectively combines information about two time points – CRISPR barcoding during early development, and trajectory inference at the time of analysis – and can serve as a blueprint for understanding the origin of transient cell types that are generated in disease conditions. With this strategy we could classify cardiac fibroblasts into two groups, based on their origin from either the epicardium or the endocardium. While the existence of epicardial fibroblasts is firmly established, we now show that there are three types of endocardial fibroblasts (*nppc* fibroblasts, *spock3* fibroblasts, valve fibroblasts). As an example for the type of high-resolution cellular and molecular dissection of regulatory interactions that becomes possible with our data, we performed an analysis of the effects of a broad perturbation, Wnt inhibition, which revealed that Wnt is required for activation of all endocardial fibroblasts.

Among the three types of endocardial fibroblasts, the *nppc* fibroblasts are unique because they are completely absent in the healthy heart and are generated in the injury area only upon deep injury directly affecting the endocardium. The epicardium-derived transient *col11/12* fibroblasts are the cell type that corresponds most closely to the “activated” fibroblasts previously described in the literature (*14*). Interestingly, they appear to be induced via retinoic acid, which is produced in a variety of cell types, including the epicardium and *nppc* fibroblasts. Hence, the described pro-regenerative function of retinoic acid is at least partially indirect, acting via induction of *col11/col12* fibroblasts, which in turn produce pro-regenerative factors. The perivascular cells are another interesting cell type with fibroblast characteristics: In contrast to previous reports, our data shows that the perivascular cells are not the only source of *nrg1* (*26*), but we found that they are the source of *cxcl12b*, a major factor in revascularization after heart injury, whose source had previously not been reported (*33*). In summary, our detailed cellular dissection of the fibrotic response in the zebrafish heart shows previously unknown sources and mechanisms of pro-regenerative programs.

This large diversity of different fibroblast cell populations challenges the concept of the fibroblast as a well-defined cell type, and suggests that fibrosis should be seen as a collective phenomenon to which a variety of cell types contribute. We here focused our analysis on cell types that have a clear pro-regenerative function based on their expression profile. Understanding the role of cell types with a less obvious function in heart regeneration, including other fibroblast populations or immune cell types, requires targeted cell type depletion experiments (*40*), which can be set up using our atlas. Only relatively few factors with a clear pro-regenerative function in heart regeneration have been discovered (*2*). It is likely that the transient cell types we identified express additional pro-regenerative genes, and our list of differentially expressed secreted factors is a powerful resource for identification of those factors.

The zebrafish heart is highly similar to the human heart, and the pathways that contribute to the regenerative capacity of the zebrafish heart are conserved. Thus, our analysis may lead us to better understand the limited regenerative capacity of the human heart, and it opens up an exciting strategy to identify novel therapeutic approaches. Our dataset, as well as the experimental and computational approaches presented here, will serve as a powerful resource for identification of candidate interactions between cell types involved in heart regeneration.

## Author contributions

B.H., S.L., B.S., D.P. and J.P.J. conceived and designed the project. B.H. performed single-cell experiments and mRNA and cell type analysis, with support by B.S.. S.L. and M.G.S. did cryoinjury experiments. S.L. performed and analyzed histology, RNAscope, CreLox lineage tracing and Wnt inhibition experiments. B.S. built single-cell lineage trees, developed tree analysis methods, and performed trajectory and RNA velocity analysis. H.A. adapted bulk deconvolution to tomo-seq data, with guidance from B.S. and F.T.. D.P. and J.P.J. guided experiments, and J.P.J. guided data analysis. B.H., B.S. and J.P.J. wrote the manuscript, in close interaction with S.L. and D.P., and with input from all other authors. All authors discussed and interpreted results.

## Acknowledgements

We acknowledge support by MDC/BIMSB core facilities (zebrafish, light microscopy, genomics, bioinformatics), and we thank J. Richter for help with zebrafish experiments. Work in D.P.’s laboratory was supported by the DZHK (German Centre for Cardiovascular Research) and by the BMBF (German Ministry of Education and Research). Work in J.P.J.’s laboratory was funded by a European Research Council Starting Grant (ERC-StG 715361 SPACEVAR), a Fondation Leducq Transatlantic Networks Grant (16CVD03), and a Helmholtz Incubator grant (Sparse2Big ZT-I-0007). B.H. was supported by a PhD fellowship from Studienstiftung des deutschen Volkes.

## References

1. K. D. Poss, Heart Regeneration in Zebrafish. Science. 298, 2188–2190 (2002).

2. J. M. Gonzalez-Rosa, C. E. Burns, C. G. Burns, Zebrafish heart regeneration: 15 years of discoveries. Regeneration. 4, 105–123 (2017).

3. J. M. Gonzalez-Rosa, V. Martin, M. Peralta, M. Torres, N. Mercader, Extensive scar formation and regression during heart regeneration after cryoinjury in zebrafish. Development. 138, 1663–1674 (2011).

4. K. Kikuchi, J. E. Holdway, A. A. Werdich, R. M. Anderson, Y. Fang, G. F. Egnaczyk, T. Evans, C. A. MacRae, D. Y. R. Stainier, K. D. Poss, Primary contribution to zebrafish heart regeneration by gata4+ cardiomyocytes. Nature. 464, 601–605 (2010).

5. C. Jopling, E. Sleep, M. Raya, M. Martí, A. Raya, J. C. I. Belmonte, Zebrafish heart regeneration occurs by cardiomyocyte dedifferentiation and proliferation. Nature. 464, 606–609 (2010).

6. K. Kikuchi, J. E. Holdway, R. J. Major, N. Blum, R. D. Dahn, G. Begemann, K. D. Poss, Retinoic Acid Production by Endocardium and Epicardium Is an Injury Response Essential for Zebrafish Heart Regeneration. Developmental Cell. 20, 397–404 (2011).

7. Y. Fang, V. Gupta, R. Karra, J. E. Holdway, K. Kikuchi, K. D. Poss, Translational profiling of cardiomyocytes identifies an early Jak1/Stat3 injury response required for zebrafish heart regeneration. Proceedings of the National Academy of Sciences. 110, 13416–13421 (2013).

8. Y. Han, A. Chen, K.-B. Umansky, K. A. Oonk, W.-Y. Choi, A. L. Dickson, J. Ou, V. Cigliola, O. Yifa, J. Cao, V. A. Tornini, B. D. Cox, E. Tzahor, K. D. Poss, Vitamin D Stimulates Cardiomyocyte Proliferation and Controls Organ Size and Regeneration in Zebrafish. Developmental Cell. 48, 853-863.e5 (2019).

9. C.-C. Wu, F. Kruse, M. D. Vasudevarao, J. P. Junker, D. C. Zebrowski, K. Fischer, E. S. Noël, D. Grün, E. Berezikov, F. B. Engel, A. van Oudenaarden, G. Weidinger, J. Bakkers, Spatially Resolved Genome-wide Transcriptional Profiling Identifies BMP Signaling as Essential Regulator of Zebrafish Cardiomyocyte Regeneration. Developmental Cell. 36, 36–49 (2016).

10. A. Lepilina, A. N. Coon, K. Kikuchi, J. E. Holdway, R. W. Roberts, C. G. Burns, K. D. Poss, A Dynamic Epicardial Injury Response Supports Progenitor Cell Activity during Zebrafish Heart Regeneration. Cell. 127, 607–619 (2006).

11. J. Münch, D. Grivas, Á. González-Rajal, R. Torregrosa-Carrión, J. L. de la Pompa, Notch signalling restricts inflammation and serpine1 expression in the dynamic endocardium of the regenerating zebrafish heart. Development. 144, 1425–1440 (2017).

12. A.-S. de Preux Charles, T. Bise, F. Baier, J. Marro, A. Jaźwińska, Distinct effects of inflammation on preconditioning and regeneration of the adult zebrafish heart. Open Biol. 6, 160102 (2016).

13. S.-L. Lai, R. Marín-Juez, P. L. Moura, C. Kuenne, J. K. H. Lai, A. T. Tsedeke, S. Guenther, M. Looso, D. Y. Stainier, Reciprocal analyses in zebrafish and medaka reveal that harnessing the immune response promotes cardiac regeneration. eLife. 6, e25605 (2017).

14. H. Sánchez-Iranzo, M. Galardi-Castilla, A. Sanz-Morejón, J. M. González-Rosa, R. Costa, A. Ernst, J. Sainz de Aja, X. Langa, N. Mercader, Transient fibrosis resolves via fibroblast inactivation in the regenerating zebrafish heart. Proc Natl Acad Sci USA. 115, 4188–4193 (2018).

15. 15. G. Ozhan, G. Weidinger, Cell Regeneration (2015), doi:10.1186/s13619-015-0017-8.

16. D. T. Paik, S. Cho, L. Tian, H. Y. Chang, J. C. Wu, Single-cell RNA sequencing in cardiovascular development, disease and medicine. Nat Rev Cardiol (2020), doi:10.1038/s41569-020-0359-y.

17. N. R. Tucker, M. Chaffin, S. J. Fleming, A. W. Hall, V. A. Parsons, K. C. Bedi Jr, A.-D. Akkad, C. N. Herndon, A. Arduini, I. Papangeli, C. Roselli, F. Aguet, S. H. Choi, K. G. Ardlie, M. Babadi, K. B. Margulies, C. M. Stegmann, P. T. Ellinor, Circulation (2020), doi:10.1161/CIRCULATIONAHA.119.045401.

18. M. Litviňuková, C. Talavera-López, H. Maatz, D. Reichart, C. L. Worth, E. L. Lindberg, M. Kanda, K. Polanski, M. Heinig, M. Lee, E. R. Nadelmann, K. Roberts, L. Tuck, E. S. Fasouli, D. M. DeLaughter, B. McDonough, H. Wakimoto, J. M. Gorham, S. Samari, K. T. Mahbubani, K. Saeb-Parsy, G. Patone, J. J. Boyle, H. Zhang, H. Zhang, A. Viveiros, G. Y. Oudit, O. Bayraktar, J. G. Seidman, C. E. Seidman, M. Noseda, N. Hubner, S. A. Teichmann, Cells of the adult human heart. Nature (2020), doi:10.1038/s41586-020-2797-4.

19. N. Farbehi, R. Patrick, A. Dorison, M. Xaymardan, V. Janbandhu, K. Wystub-Lis, J. W. Ho, R. E. Nordon, R. P. Harvey, Single-cell expression profiling reveals dynamic flux of cardiac stromal, vascular and immune cells in health and injury. eLife. 8, e43882 (2019).

20. H. Honkoop, D. E. de Bakker, A. Aharonov, F. Kruse, A. Shakked, P. D. Nguyen, C. de Heus, L. Garric, M. J. Muraro, A. Shoffner, F. Tessadori, J. C. Peterson, W. Noort, A. Bertozzi, G. Weidinger, G. Posthuma, D. Grün, W. J. van der Laarse, J. Klumperman, R. T. Jaspers, K. D. Poss, A. van Oudenaarden, E. Tzahor, J. Bakkers, Single-cell analysis uncovers that metabolic reprogramming by ErbB2 signaling is essential for cardiomyocyte proliferation in the regenerating heart. eLife. 8, e50163 (2019).

21. B. Spanjaard, B. Hu, N. Mitic, P. Olivares-Chauvet, S. Janjuha, N. Ninov, J. P. Junker, Simultaneous lineage tracing and cell-type identification using CRISPR-Cas9-induced genetic scars. Nature Biotech. 36, 469–473 (2018).

22. B. Raj, D. E. Wagner, A. McKenna, S. Pandey, A. M. Klein, J. Shendure, J. A. Gagnon, F. Schier, Simultaneous single-cell profiling of lineages and cell types in the vertebrate brain. Nature Biotech. 36, 442–450 (2018).

23. A. Alemany, M. Florescu, C. S. Baron, J. Peterson-Maduro, A. van Oudenaarden, Whole-organism clone tracing using single-cell sequencing. Nature. 556, 1–22 (2018).

24. J. P. Junker, E. S. Noël, V. Guryev, K. A. Peterson, G. Shah, J. Huisken, A. P. McMahon, E. Berezikov, J. Bakkers, A. van Oudenaarden, Genome-wide RNA Tomography in the Zebrafish Embryo. Cell. 159, 662–675 (2014).

25. H. Aliee, F. Theis, “AutoGeneS: Automatic gene selection using multi-objective optimization for RNA-seq deconvolution” (preprint, Bioinformatics, 2020), doi:10.1101/2020.02.21.940650.

26. M. Gemberling, R. Karra, A. L. Dickson, K. D. Poss, Nrg1 is an injury-induced cardiomyocyte mitogen for the endogenous heart regeneration program in zebrafish. eLife. 4, e05871 (2015).

27. J. Wang, R. Karra, A. L. Dickson, K. D. Poss, Fibronectin is deposited by injury-activated epicardial cells and is necessary for zebrafish heart regeneration. Developmental Biology. 382, 427–435 (2013).

28. J. Marro, C. Pfefferli, A.-S. de Preux Charles, T. Bise, A. Jaźwińska, Collagen XII Contributes to Epicardial and Connective Tissues in the Zebrafish Heart during Ontogenesis and Regeneration. PLoS ONE. 11, e0165497 (2016).

29. D. Wehner, T. M. Tsarouchas, A. Michael, C. Haase, G. Weidinger, M. M. Reimer, T. Becker, C. G. Becker, Wnt signaling controls pro-regenerative Collagen XII in functional spinal cord regeneration in zebrafish. Nat Commun. 8, 126 (2017).

30. P. Bouillet, M. Oulad-Abdelghani, S. Vicaire, J.-M. Garnier, B. Schuhbaur, P. Dollé, P. Chambon, Efficient Cloning of cDNAs of Retinoic Acid-Responsive Genes in P19 Embryonal Carcinoma Cells and Characterization of a Novel Mouse Gene, Stra1 (Mouse LERK-2/Eplg2). Developmental Biology. 170, 420–433 (1995).

31. A. Isken, M. Golczak, V. Oberhauser, S. Hunzelmann, W. Driever, Y. Imanishi, K. Palczewski, J. von Lintig, RBP4 Disrupts Vitamin A Uptake Homeostasis in a STRA6-Deficient Animal Model for Matthew-Wood Syndrome. Cell Metabolism. 7, 258–268 (2008).

32. G. Bergers, S. Song, The role of pericytes in blood-vessel formation and maintenance. Neuro-Oncology. 7, 452–464 (2005).

33. R. Marín-Juez, H. El-Sammak, C. S. M. Helker, A. Kamezaki, S. T. Mullapuli, S.-I. Bibli, M. J. Foglia, I. Fleming, K. D. Poss, D. Y. R. Stainier, Coronary Revascularization During Heart Regeneration Is Regulated by Epicardial and Endocardial Cues and Forms a Scaffold for Cardiomyocyte Repopulation. Developmental Cell. 51, 503-515.e4 (2019).

34. F. A. Wolf, F. K. Hamey, M. Plass, J. Solana, J. S. Dahlin, B. Göttgens, N. Rajewsky, L. Simon, F. J. Theis, PAGA: graph abstraction reconciles clustering with trajectory inference through a topology preserving map of single cells. Genome Biology. 20, 59 (2019).

35. G. La Manno, R. Soldatov, A. Zeisel, E. Braun, H. Hochgerner, V. Petukhov, K. Lidschreiber, M. E. Kastriti, P. Lönnerberg, A. Furlan, J. Fan, L. E. Borm, Z. Liu, D. Bruggen, J. Guo, X. He, R. Barker, E. Sundström, G. Castelo-Branco, P. Cramer, I. Adameyko, S. Linnarsson, P. V. Kharchenko, RNA velocity of single cells. Nature. 560, 1–25 (2018).

36. B. Chen, M. E. Dodge, W. Tang, J. Lu, Z. Ma, C.-W. Fan, S. Wei, W. Hao, J. Kilgore, N. S. Williams, M. G. Roth, J. F. Amatruda, C. Chen, L. Lum, Small molecule–mediated disruption of Wnt-dependent signaling in tissue regeneration and cancer. Nat Chem Biol. 5, 100–107 (2009).

37. L. Zhao, R. Ben-Yair, C. E. Burns, C. G. Burns, Endocardial Notch Signaling Promotes Cardiomyocyte Proliferation in the Regenerating Zebrafish Heart through Wnt Pathway Antagonism. Cell Reports. 26, 546-554.e5 (2019).

38. O. Aisagbonhi, M. Rai, S. Ryzhov, N. Atria, I. Feoktistov, A. K. Hatzopoulos, Experimental myocardial infarction triggers canonical Wnt signaling and endothelial-to-mesenchymal transition. Disease Models & Mechanisms. 4, 469–483 (2011).

39. J. Duan, C. Gherghe, D. Liu, E. Hamlett, L. Srikantha, L. Rodgers, J. N. Regan, M. Rojas, M. Willis, A. Leask, M. Majesky, A. Deb, Wnt1/βcatenin injury response activates the epicardium and cardiac fibroblasts to promote cardiac repair: Wnt1/βcatenin injury response regulates cardiac repair. The EMBO Journal. 31, 429–442 (2012).

40. J. R. Mathias, Z. Zhang, M. T. Saxena, J. S. Mumm, Enhanced Cell-Specific Ablation in Zebrafish Using a Triple Mutant of Escherichia ColiNitroreductase. Zebrafish. 11, 85–97 (2014).

