## Supplemental figures for "Cellular drivers of injury response and regeneration in the adult zebrafish heart"

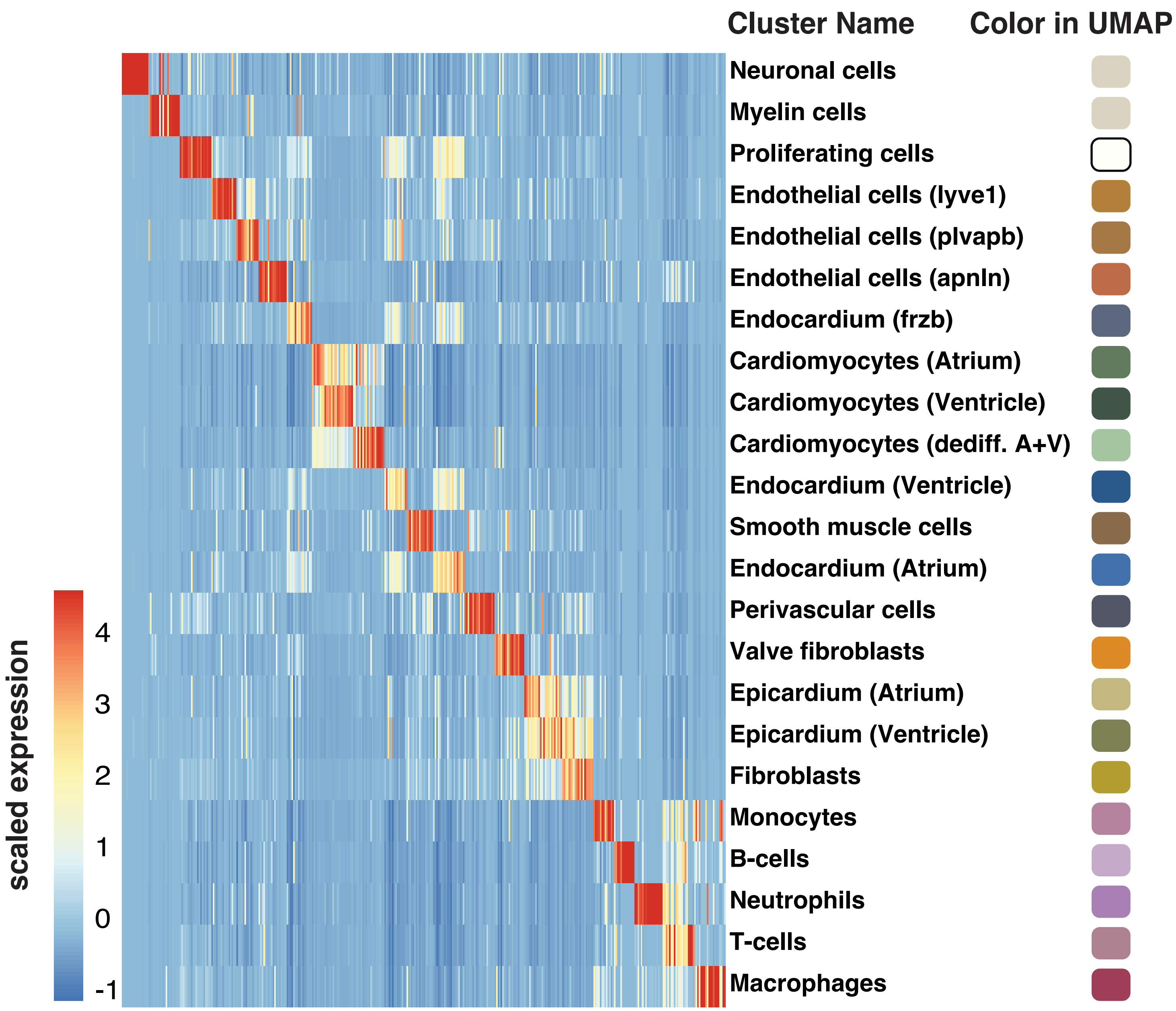


**Figure S1. Top differentially expressed genes for each cell type shown in Fig. 1B.** The genes used in this heatmap can be found in Table S2. The legend on the right shows the color of the respective cell type in Fig. 1B.

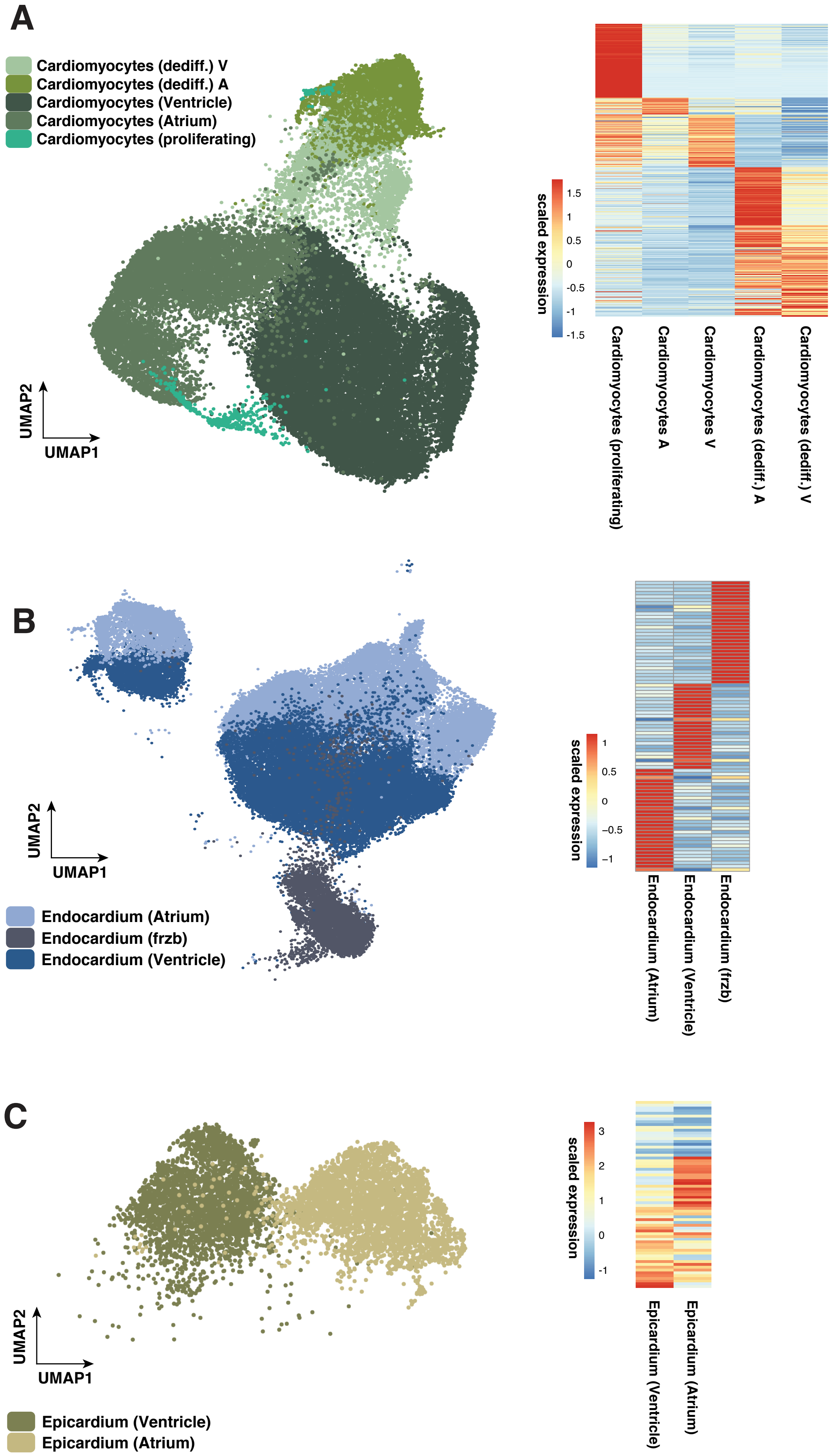


**Figure S2. Subclustering of cardiomyocytes (A), endocardium (B), and epicardium (C).** Heatmaps show top differentially expressed genes of each sub-type. The genes used for the heatmap can be found in Table S2. The UMAP representation of the clustering is zoomed in from Fig. 1B.

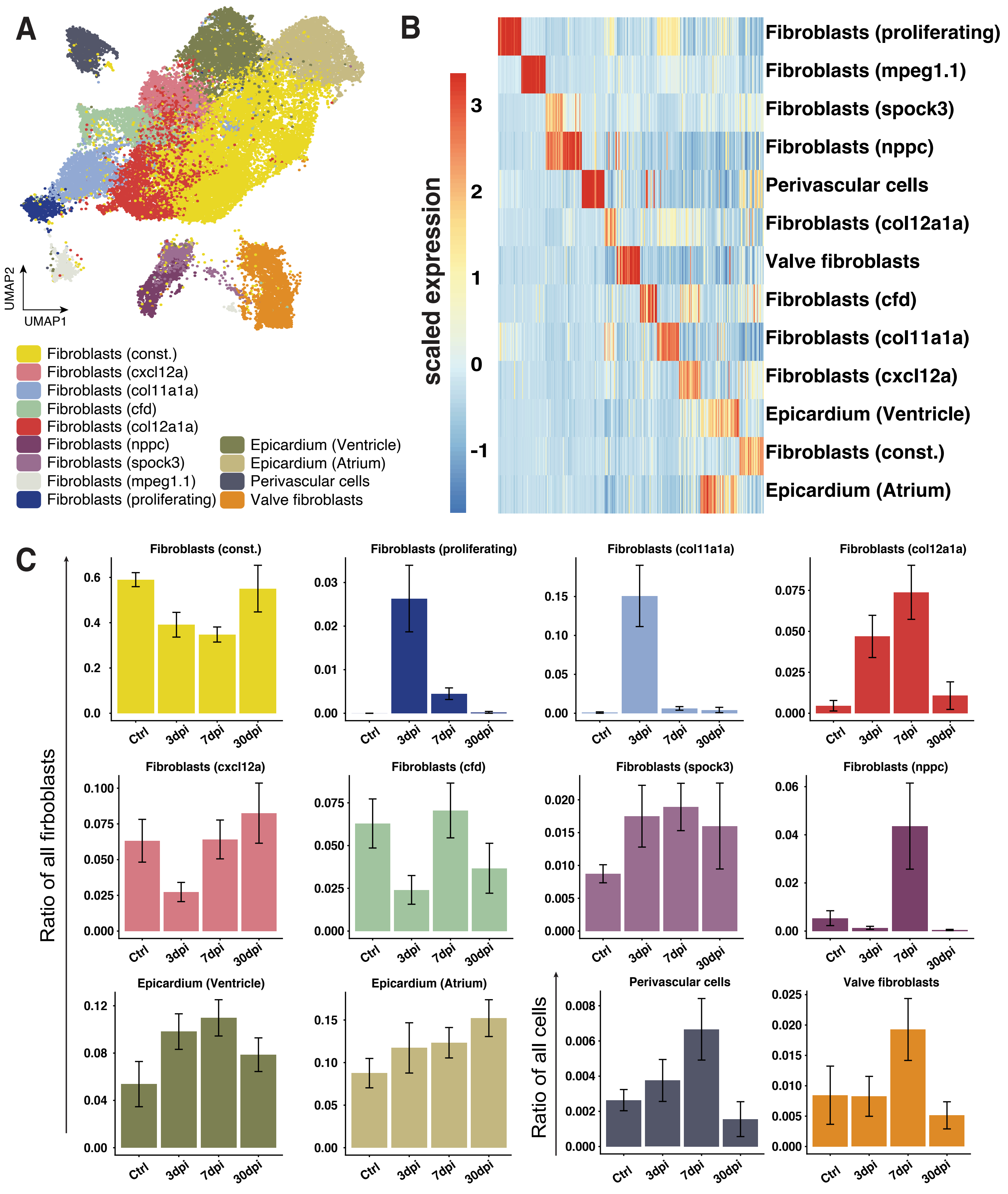


**Figure S3. Subclustering of collagen expressing cells. (A)** UMAP representation of subclustering results. Our analysis identified 13 subclusters. **(B)** Top differentially expressed genes in each subcluster. Gene names can be found in Table S3. **(C)** Cell number dynamics of all subtypes during regeneration. Mean value across all replicates of the same timepoint is shown, error bars indicate standard error of the mean.

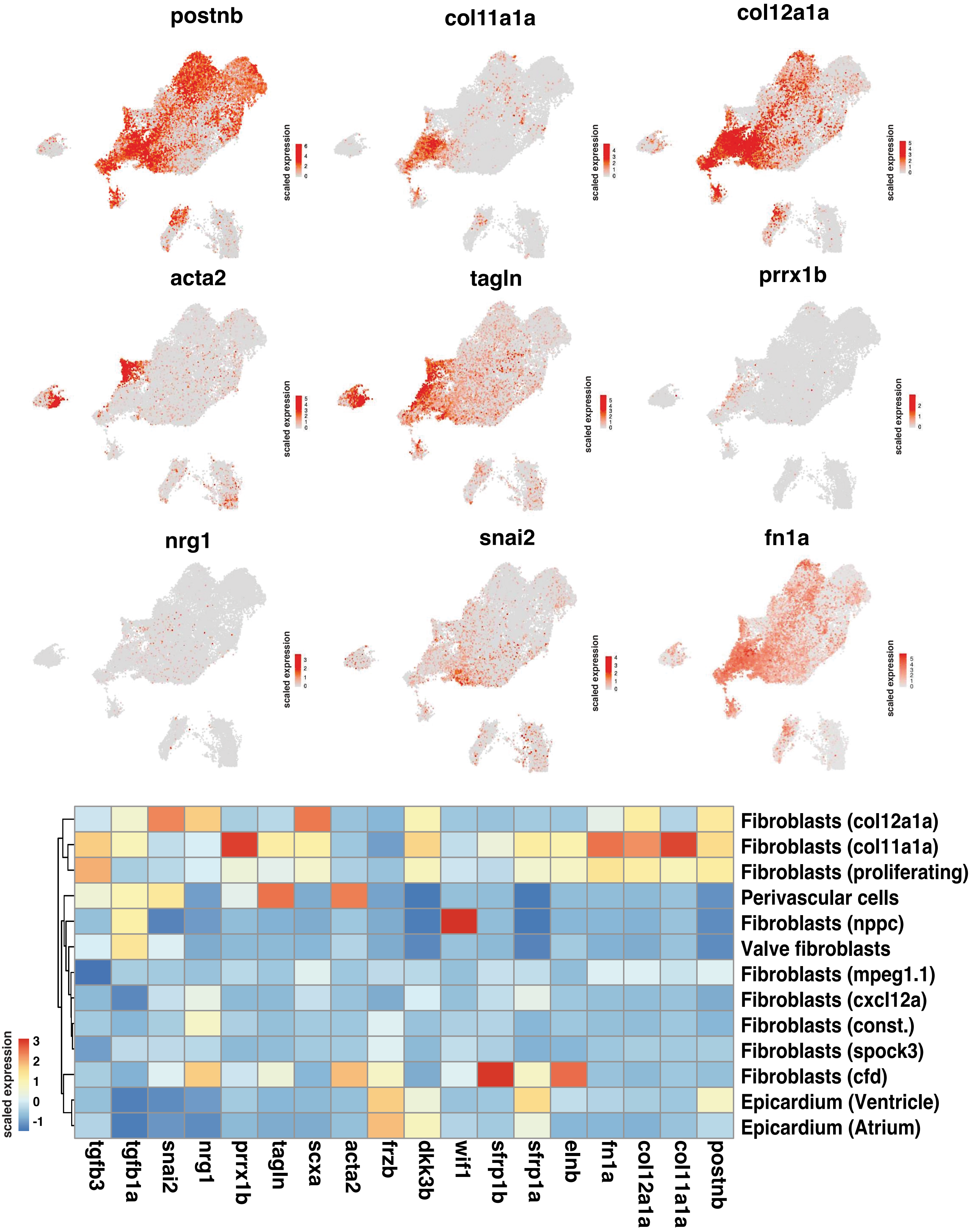


**Figure S4. Expression pattern of selected genes in fibroblast subtypes.** Expression of selected genes involved in fibroblast function or heart regeneration in the identified fibroblast subclusters.

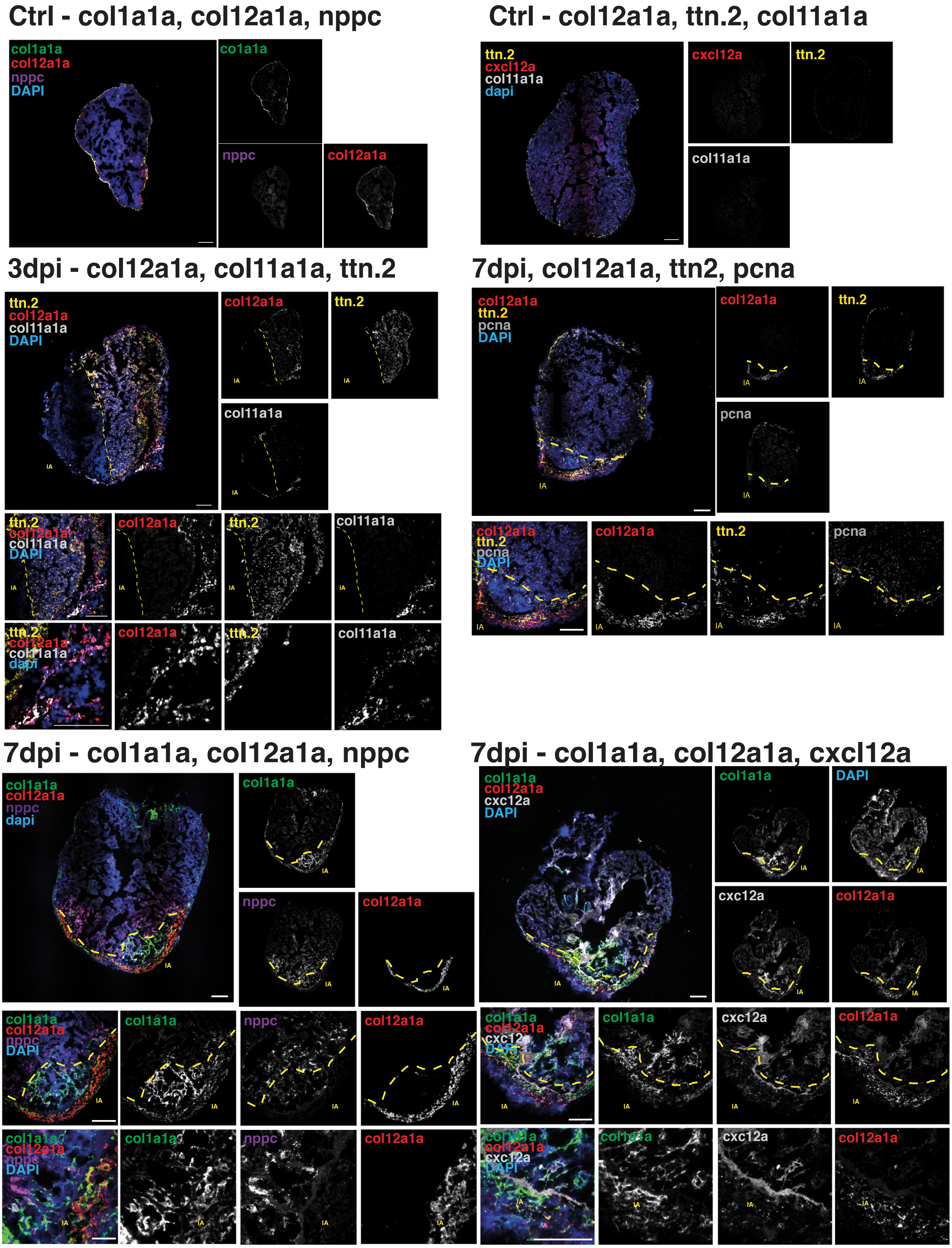


**Figure S5. Fluorescent in-situ hybridization of marker genes for cell types of interest at different time points of injury.** Markers for transient cell types (col12a1a, col11a1a, ttn.2, nppc) are localized at the injury area. Scale bar: 100$\mu$m.

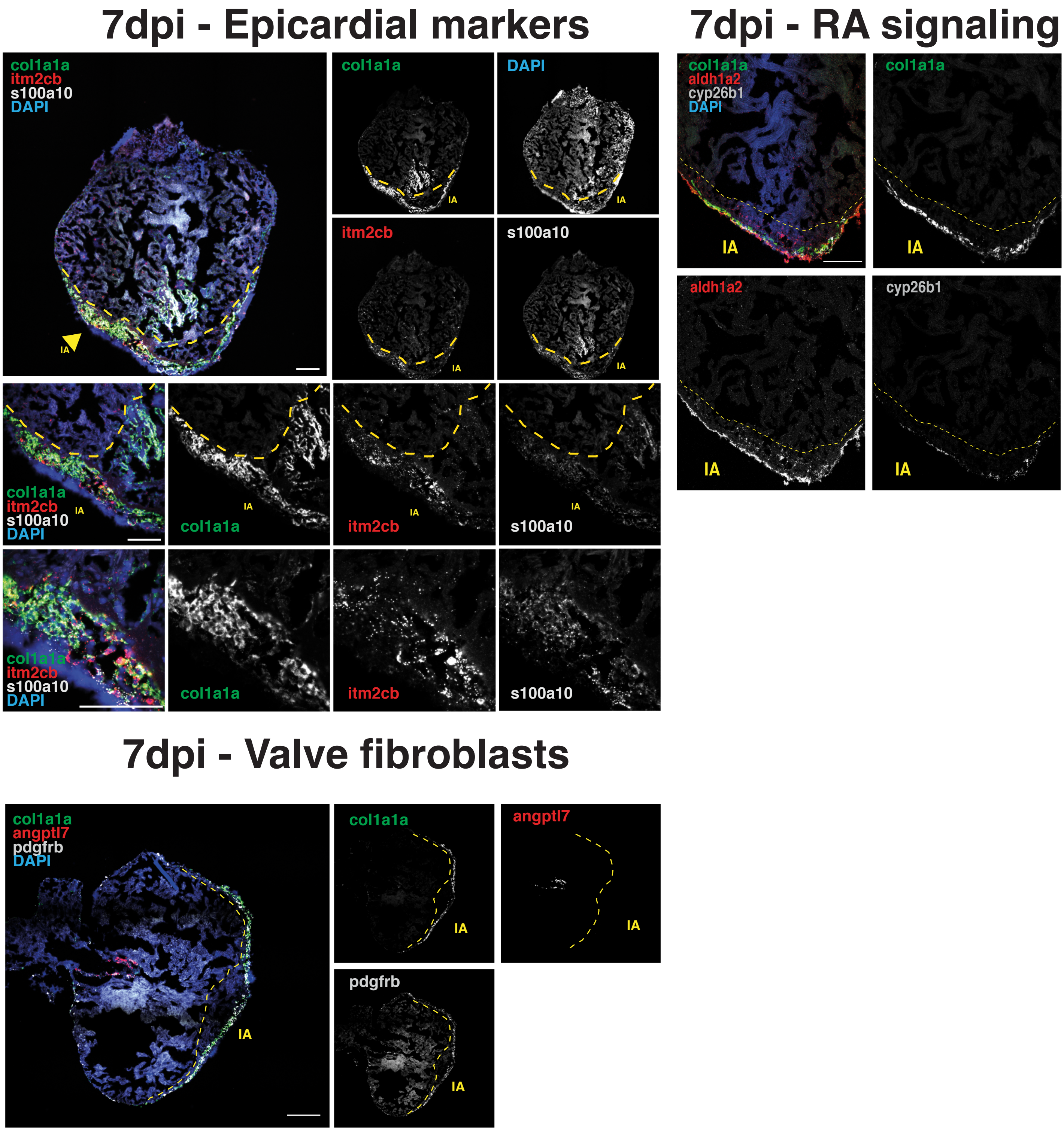


**Figure S6. Fluorescent in-situ hybridization of marker genes for additional cell types of interest and factors of the RA signaling pathway.** Epicardium – itm2cb, s100a10; valve fibroblasts – angptl7; perivascular cells – pdgfrb. Scale bar: 100$\mu$m

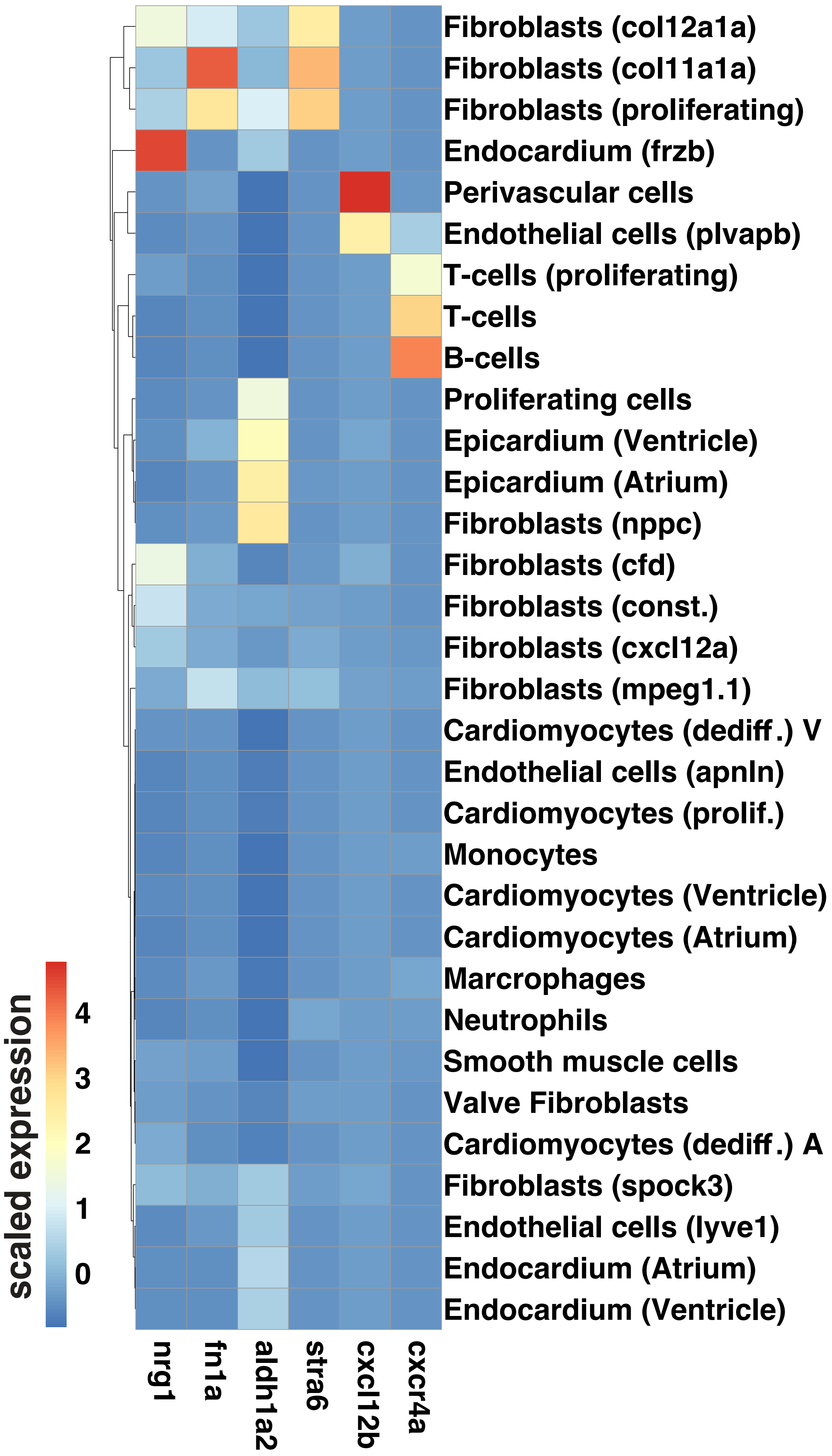


**Figure S7. Expression of known pro-regenerative factors across additional cell types of the zebrafish heart.** Expression patterns of cxcl12b, cxcr4a, aldh1a2, stra6, nrg1, and fn1a. See main text for interpretation of the expression patterns.

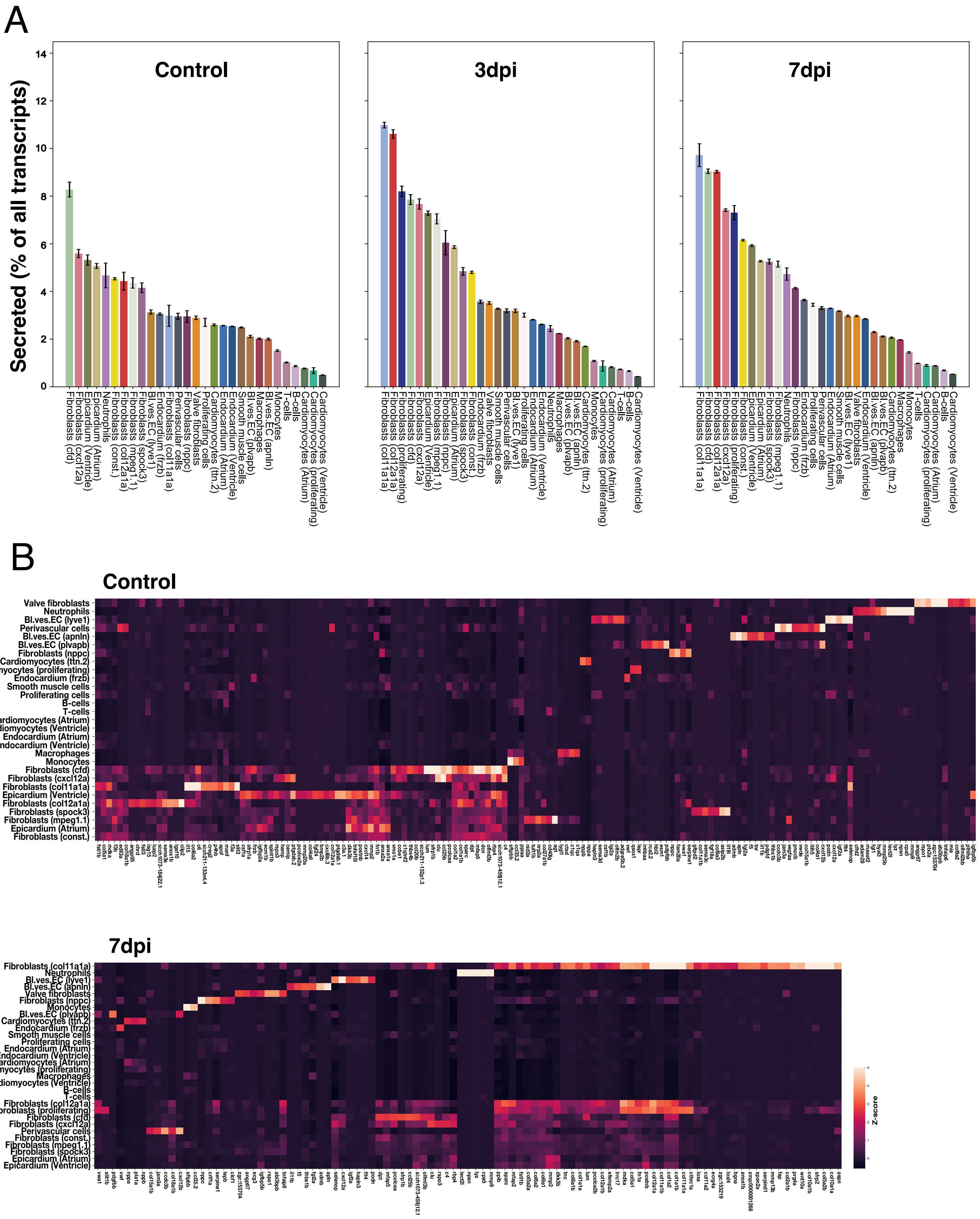


**Figure S8. Secretome analysis. (A)** All cell types, ranked by secretome expression. **(B)** Highly expressed expressed secretome genes in control hearts and at 7 dpi.

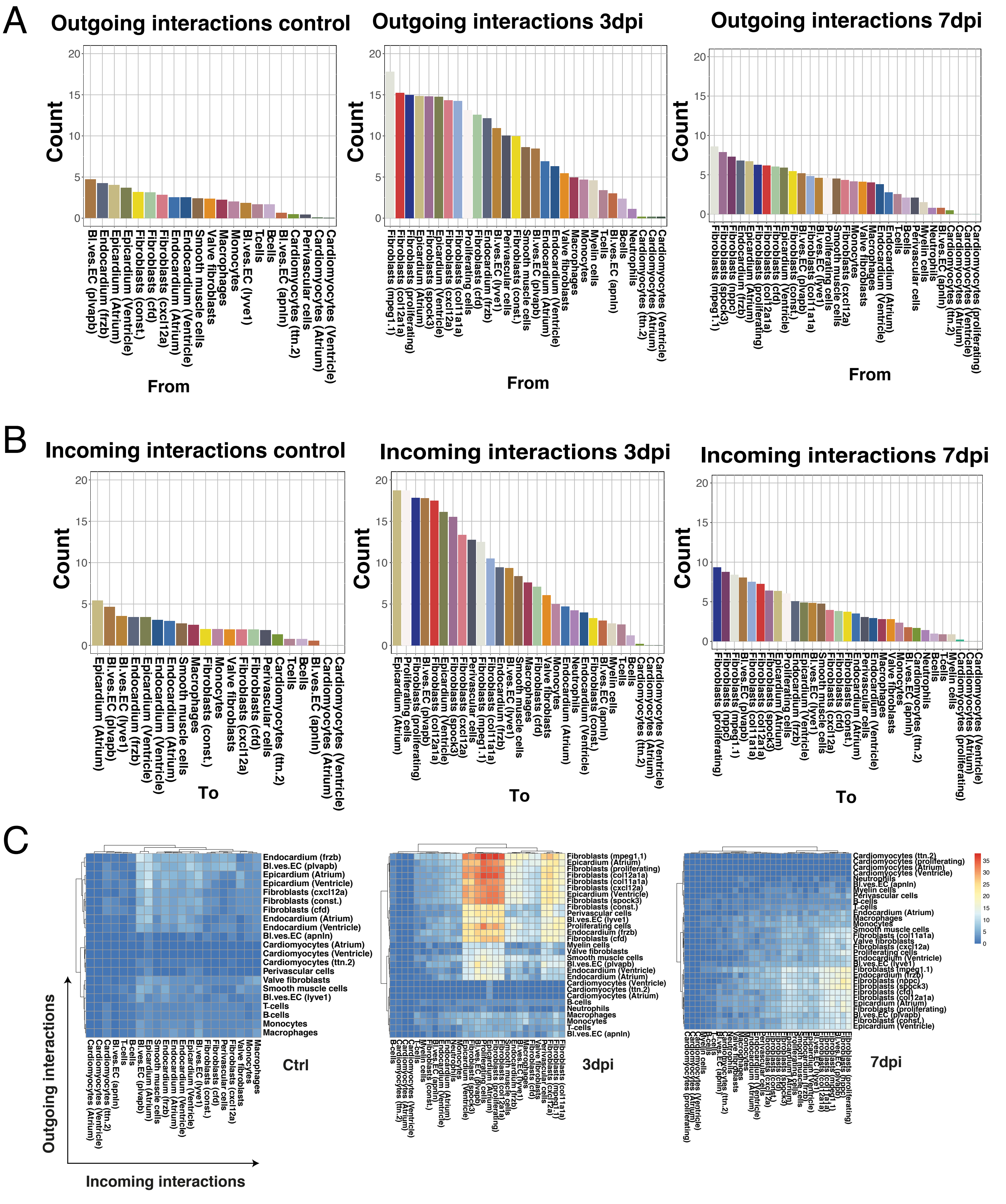


**Figure S9. Ligand-receptor analysis for cell-cell interactions. (A)** Number of outgoing interactions (ligands) per cell type. **(B)** Number of incoming interactions (receptors) per cell type. **(C)** Interactions between pairs of cell types.

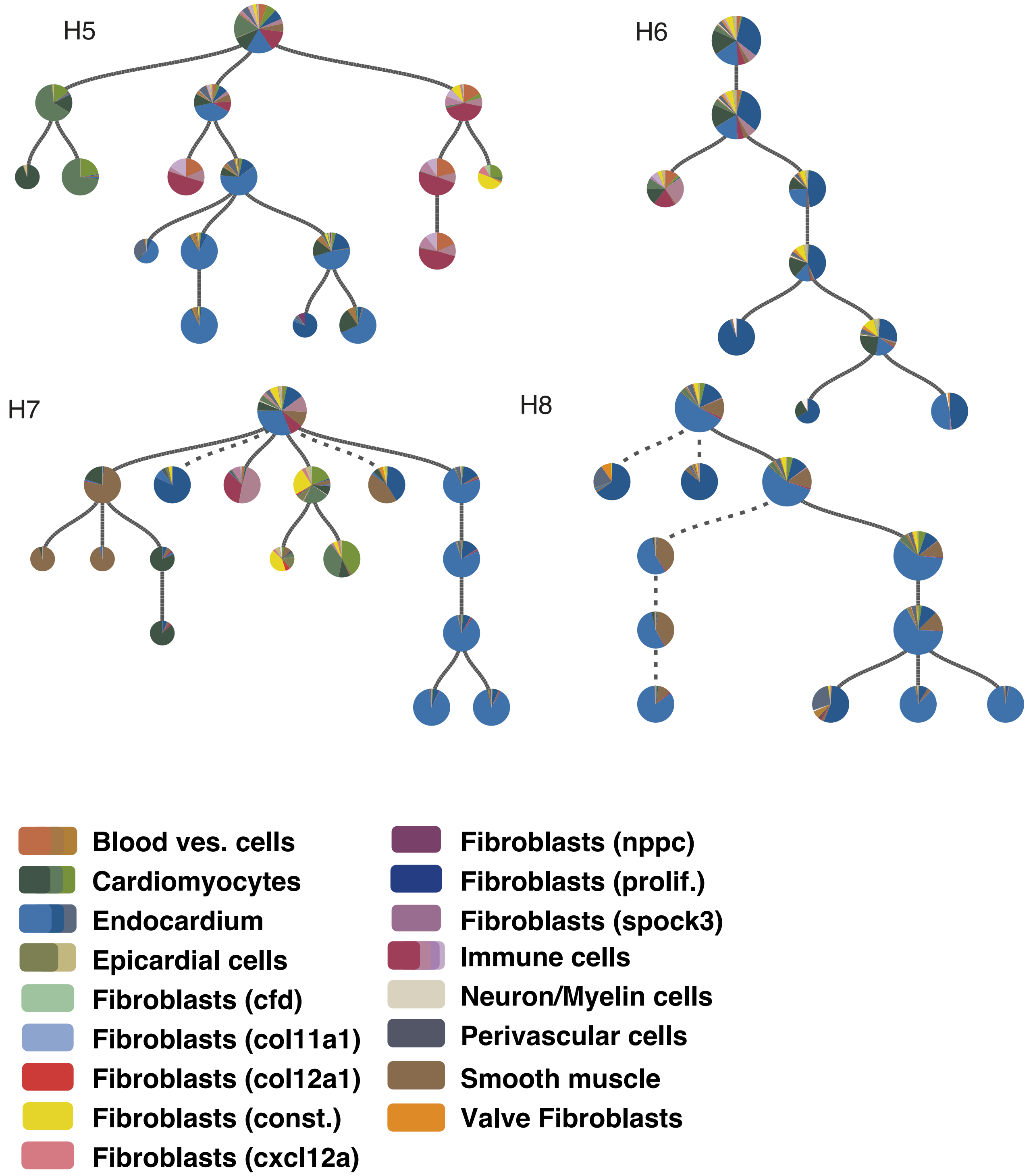


**Figure S10. Lineage trees pre-injury.**

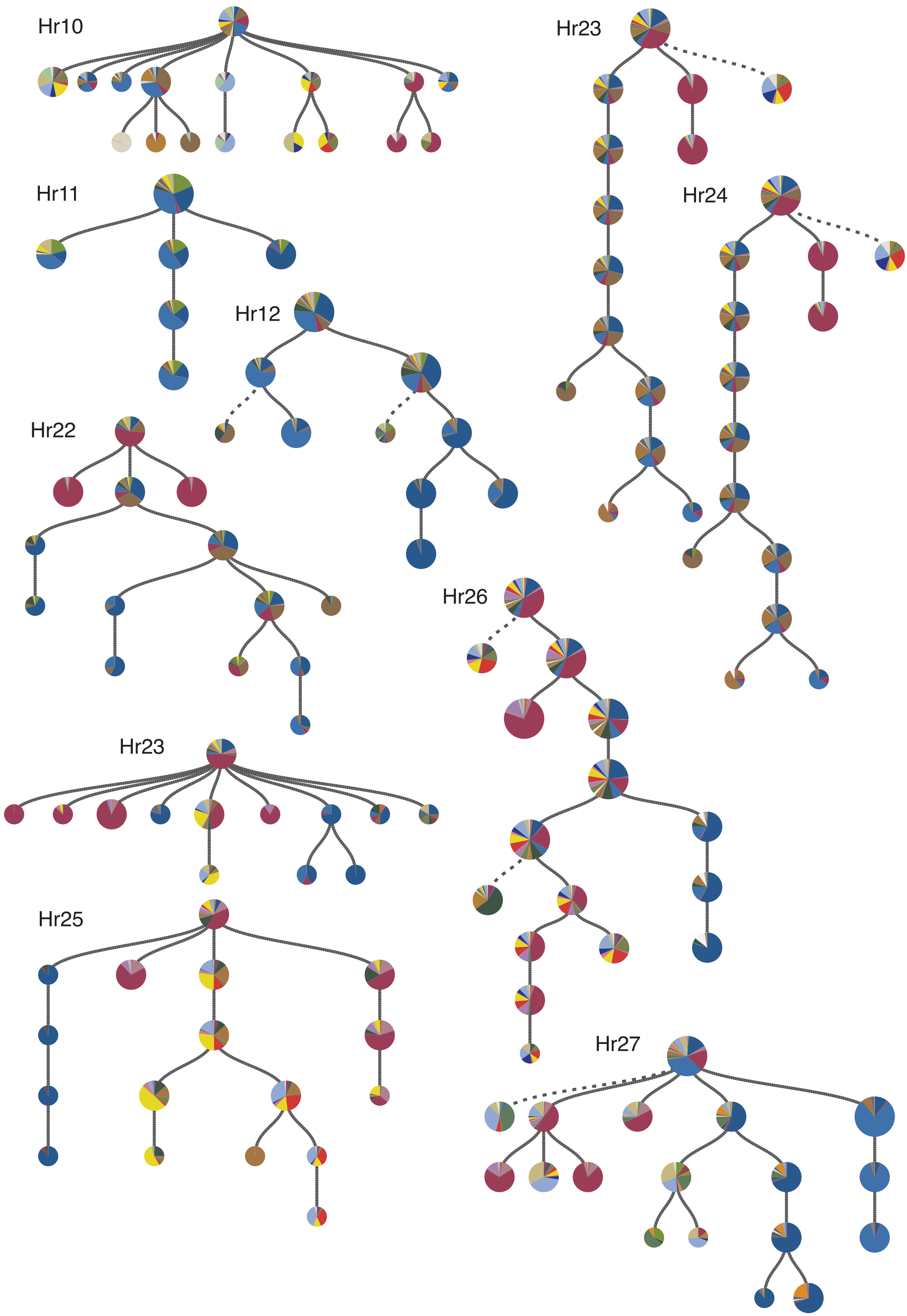


**Figure S11. Lineage trees three days post injury.** See Fig. S10 for cell type colors.

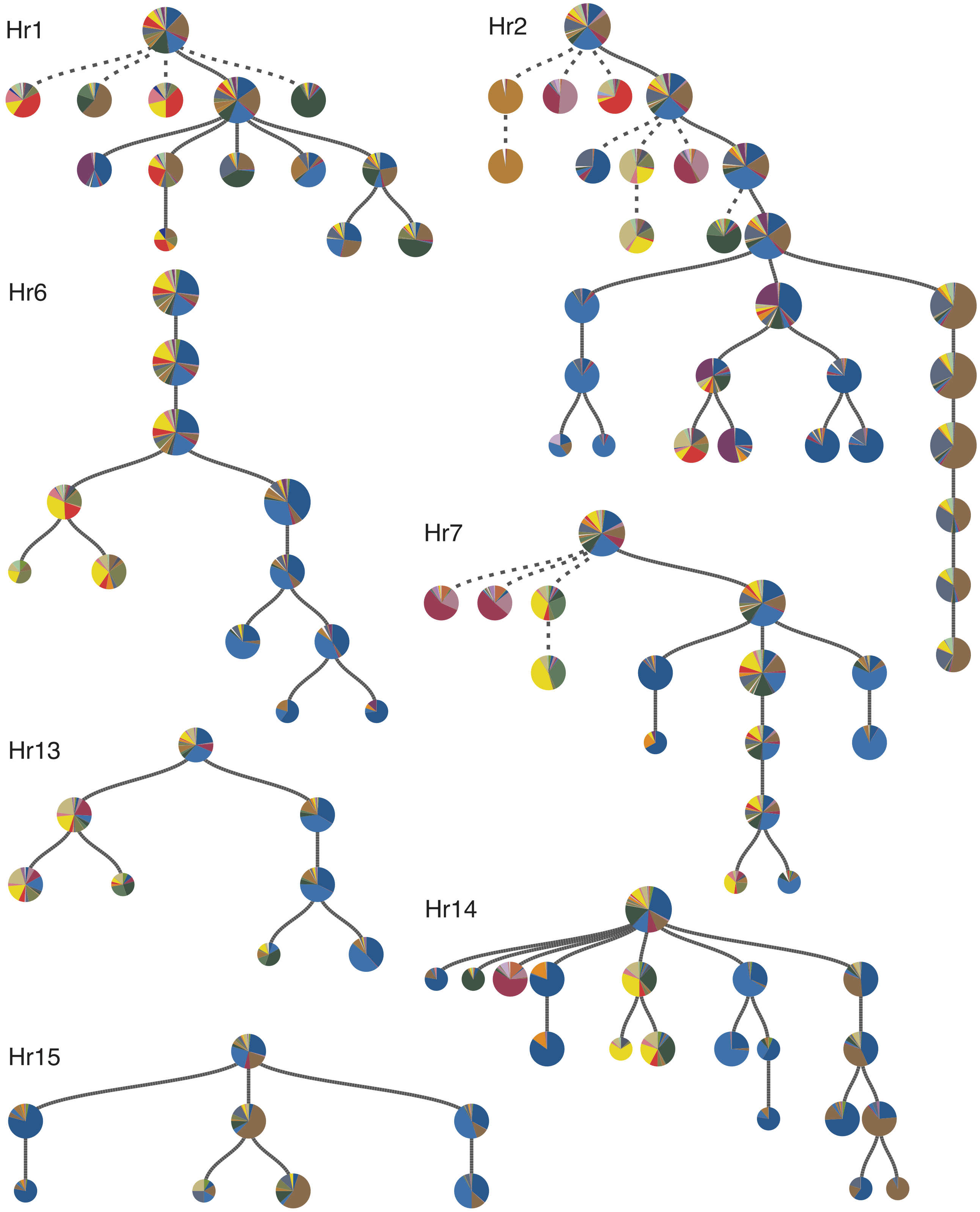


**Figure S12. Lineage trees seven days post injury.** See Fig. S10 for cell type colors.

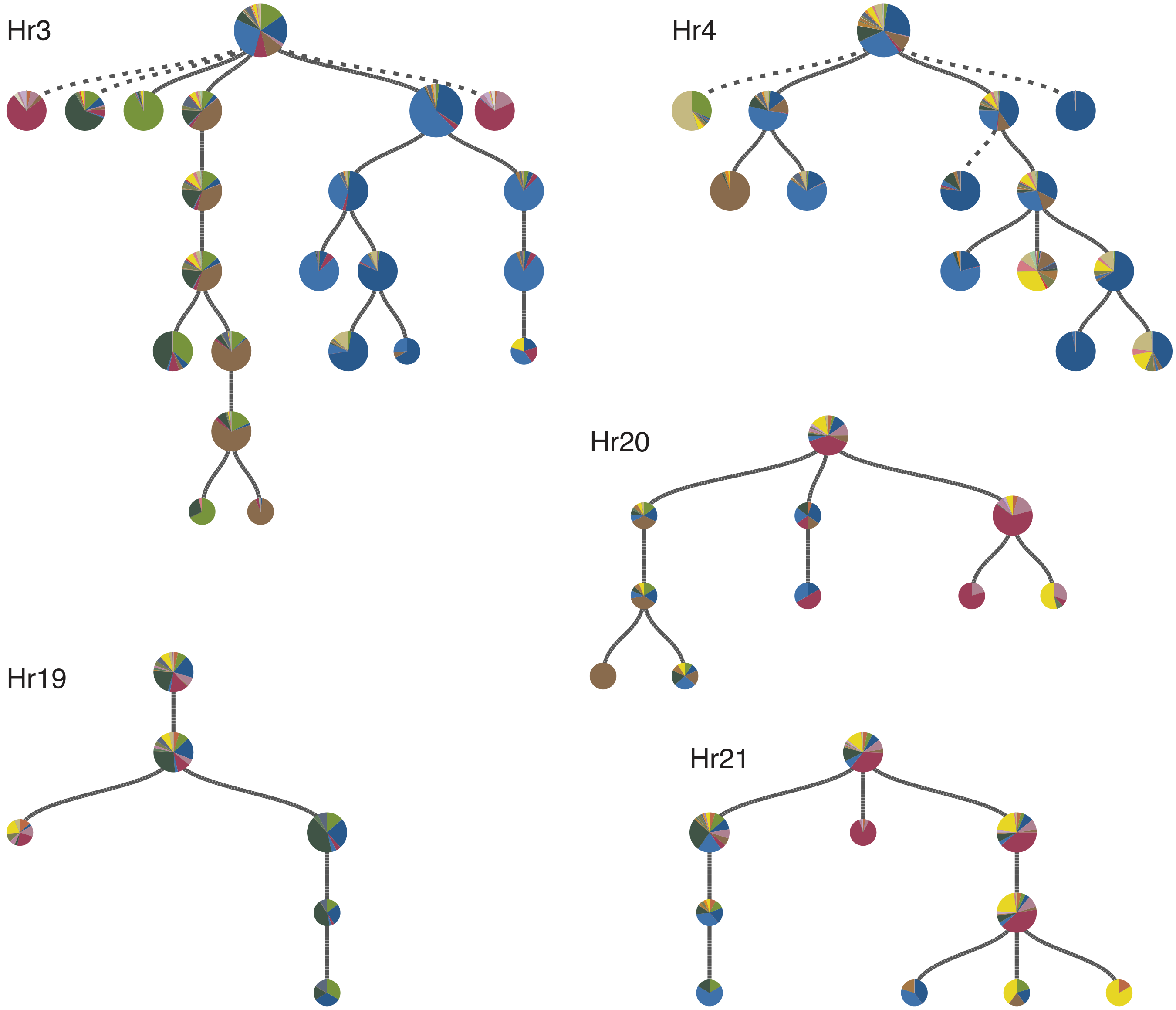


**Figure S13. Lineage trees thirty days post injury.** See Fig. S10 for cell type colors.

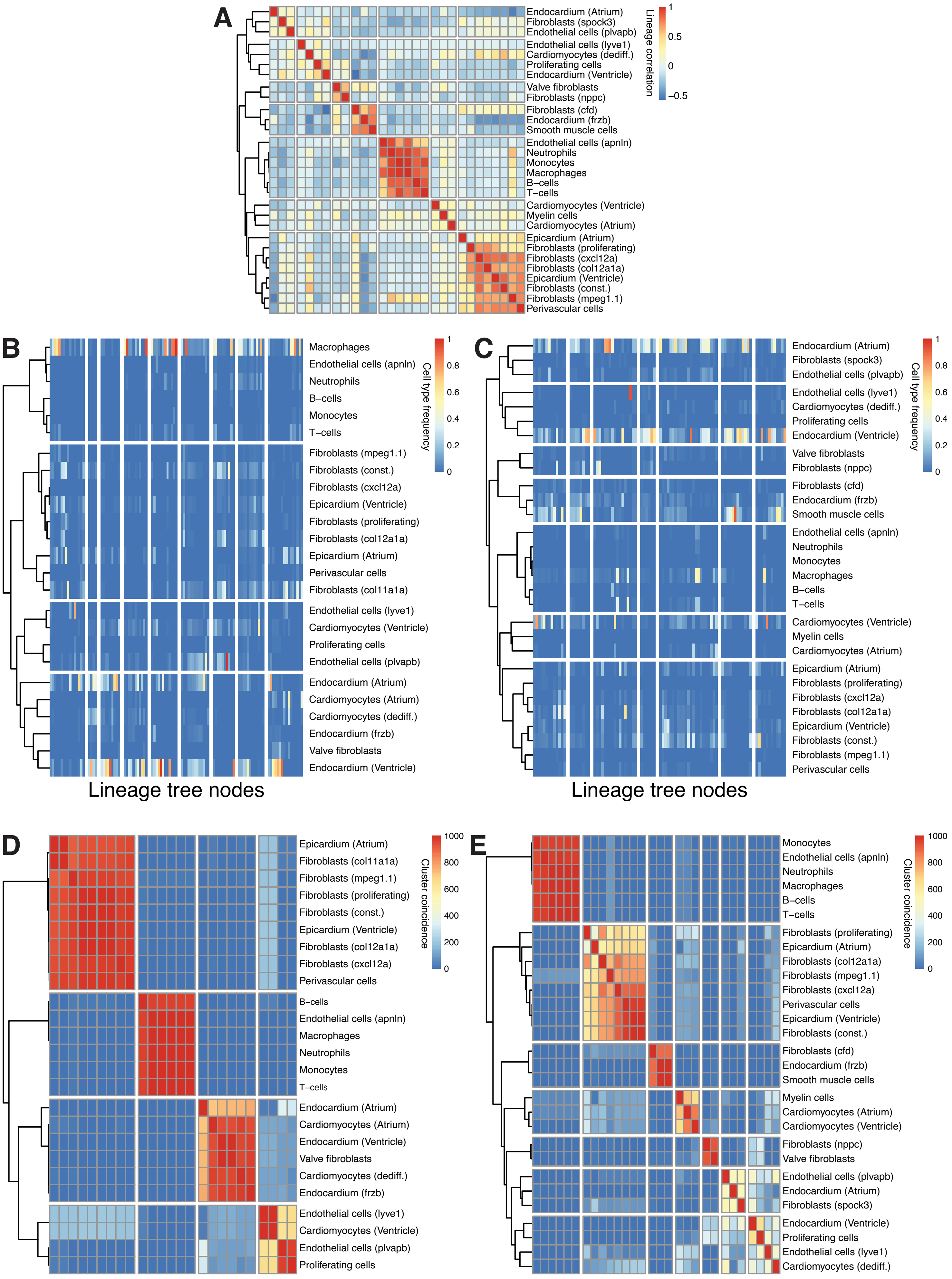


**Figure S14. Identification of lineage-related cell types at 3 dpi and 7 dpi. (A)** Cell type clustering based on lineage correlations at 7 dpi. **(B-C)** Cell type fractions per node in 3 dpi **(B)** and 7 dpi **(C)** trees. **(D-E)** Clustering at 3 dpi **(D)** and 7 dpi **(E)** is stable under 50% downsampling of trees.

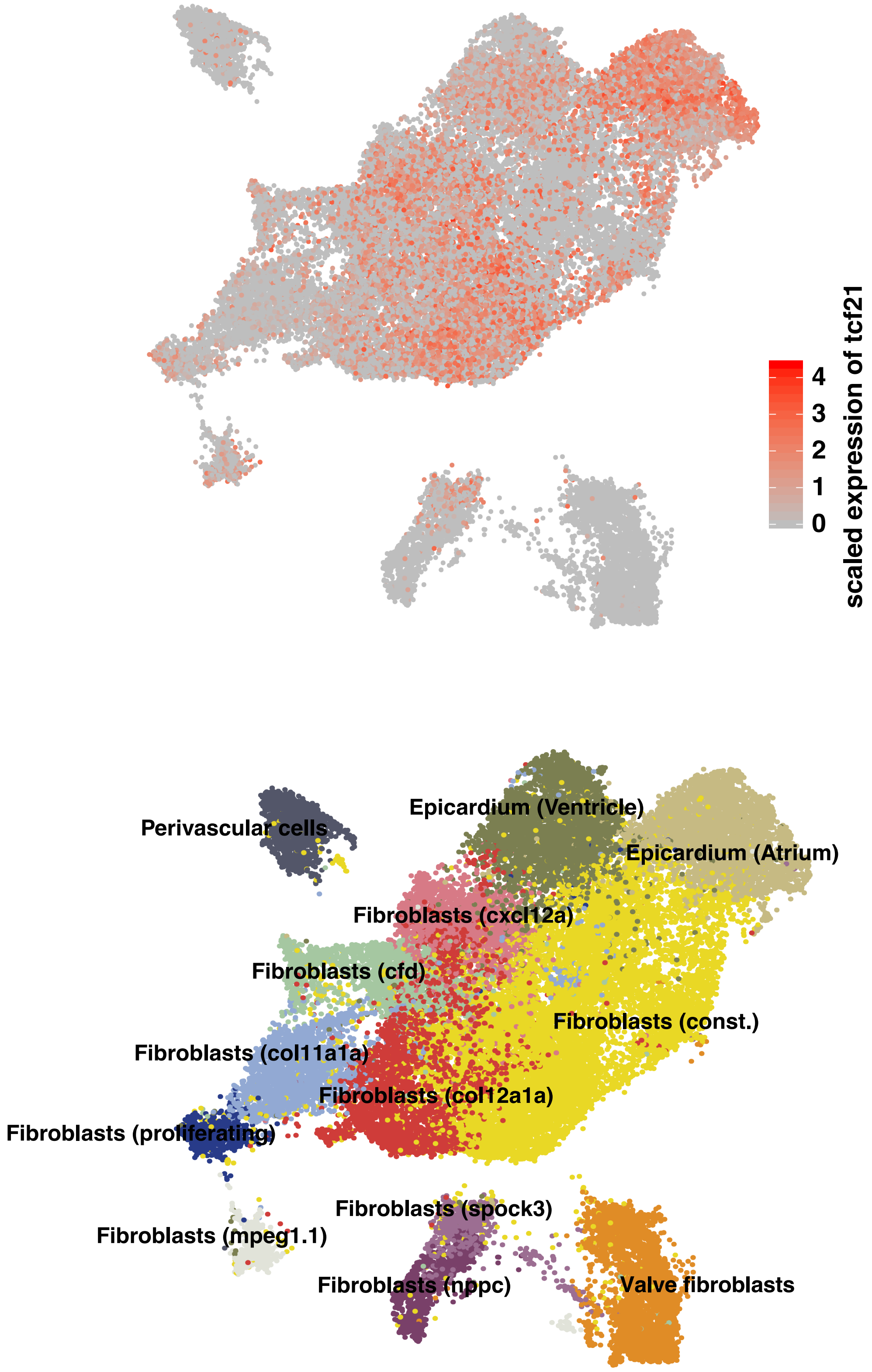


**Figure S15. Expression pattern of tcf21.** Tcf21 is expressed in epicardial cells, constitutive fibroblasts and other fibroblast types of the epicardial cluster.

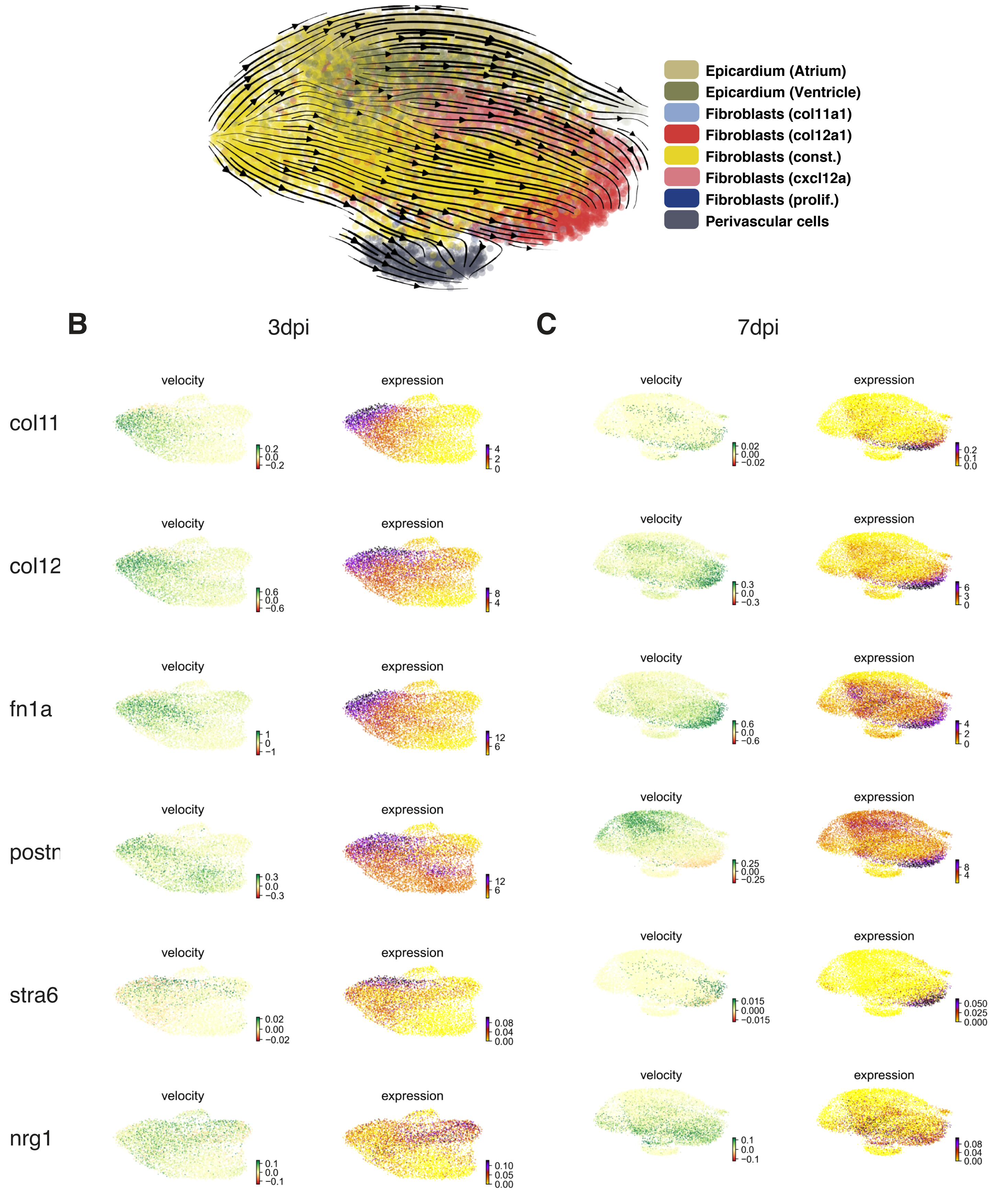


**Figure S16. RNA velocity analysis of the 3 dpi and 7 dpi epicardial niche. (A)** RNA velocity implies constitutive fibroblasts as a source of col12 fibroblasts. **(B-C)** Increase of marker genes towards niche fibroblasts at 3 dpi **(B)** and 7 dpi **(C)**.

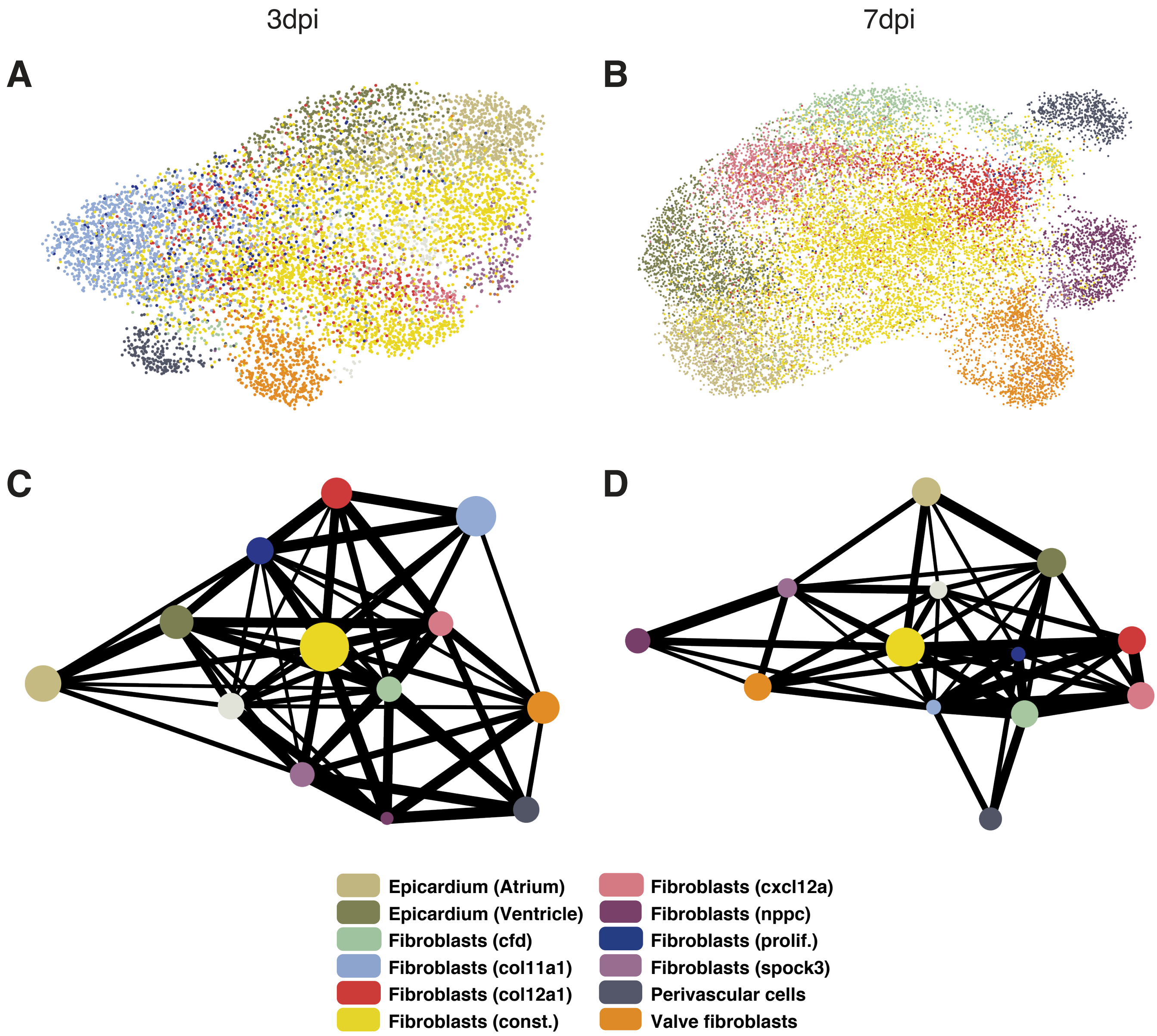


**Figure S17. Trajectory analysis on mRNA identifies connections between all niche fibroblasts. (A-B)** UMAP representation of cell types in the epicardial/fibroblast niche at 3 dpi **(A)** and 7 dpi **(B)**. **(C-D)** All cell types are connected with a strength of at least 0.3 in a PAGA analysis at 3 dpi **(C)** and 7 dpi **(D)**.

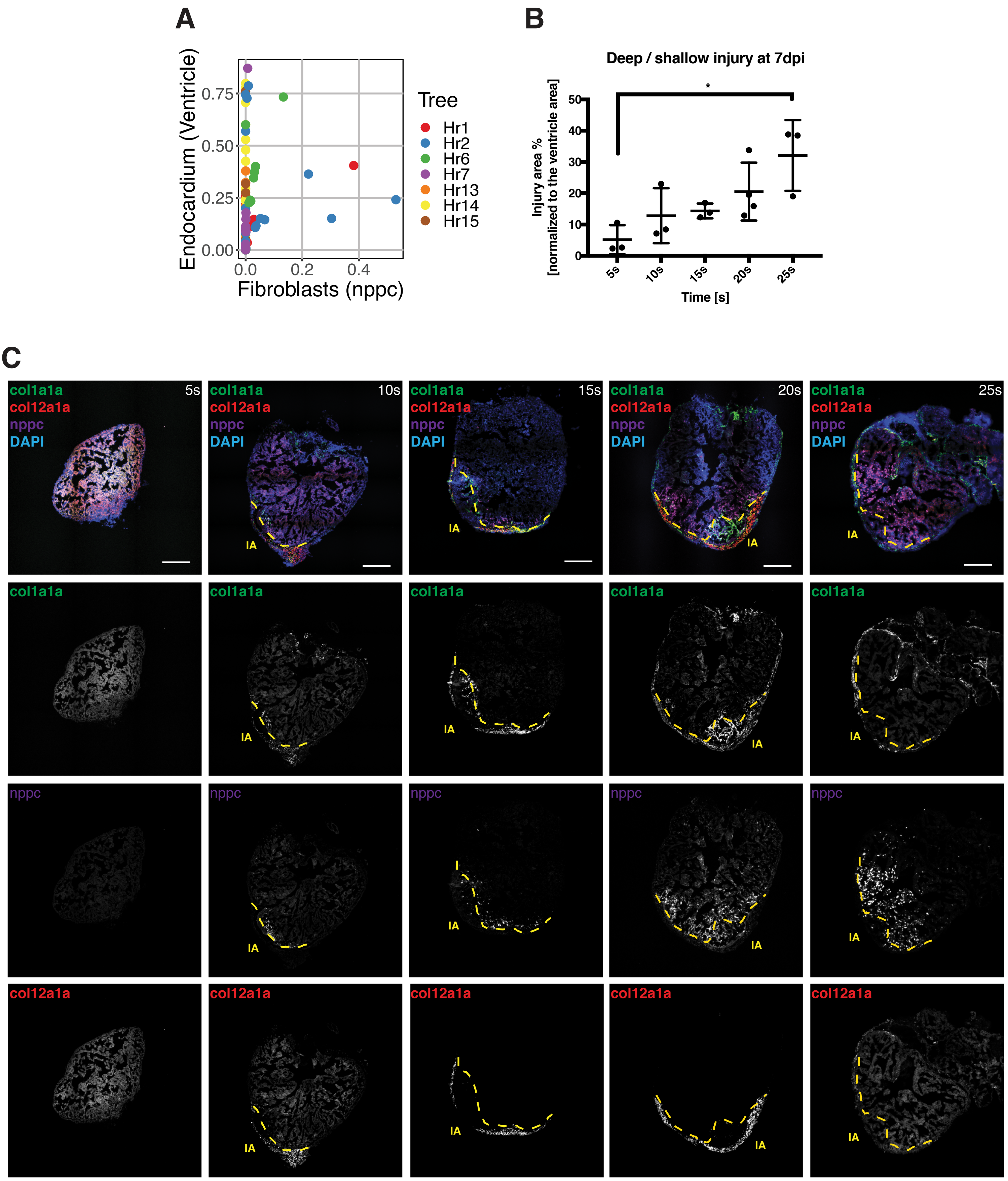


**Figure S18. Deeper injuries cause higher nppc expression. (A)** Ratio between nppc fibroblasts and ventricular endocardium in tree nodes varies per sample. **(B)** Quantification of deep and shallow injuries at 7dpi. Injury areas (IA) in % were normalized to the ventricular area. ($n_{5s}$=3, $n_{10s}$=3, $n_{15s}$=3, $n_{20s}$=4, $n_{25s}$=3). Mean ± SD. *One-way ANOVA with Tukey's multiple comparison test. *P= 0.0128. **(C)** Injuries of 20s and 25s penetrate the heart and generate nppc expression beyond the injury area. Scale bar: 100 $\mu$m.

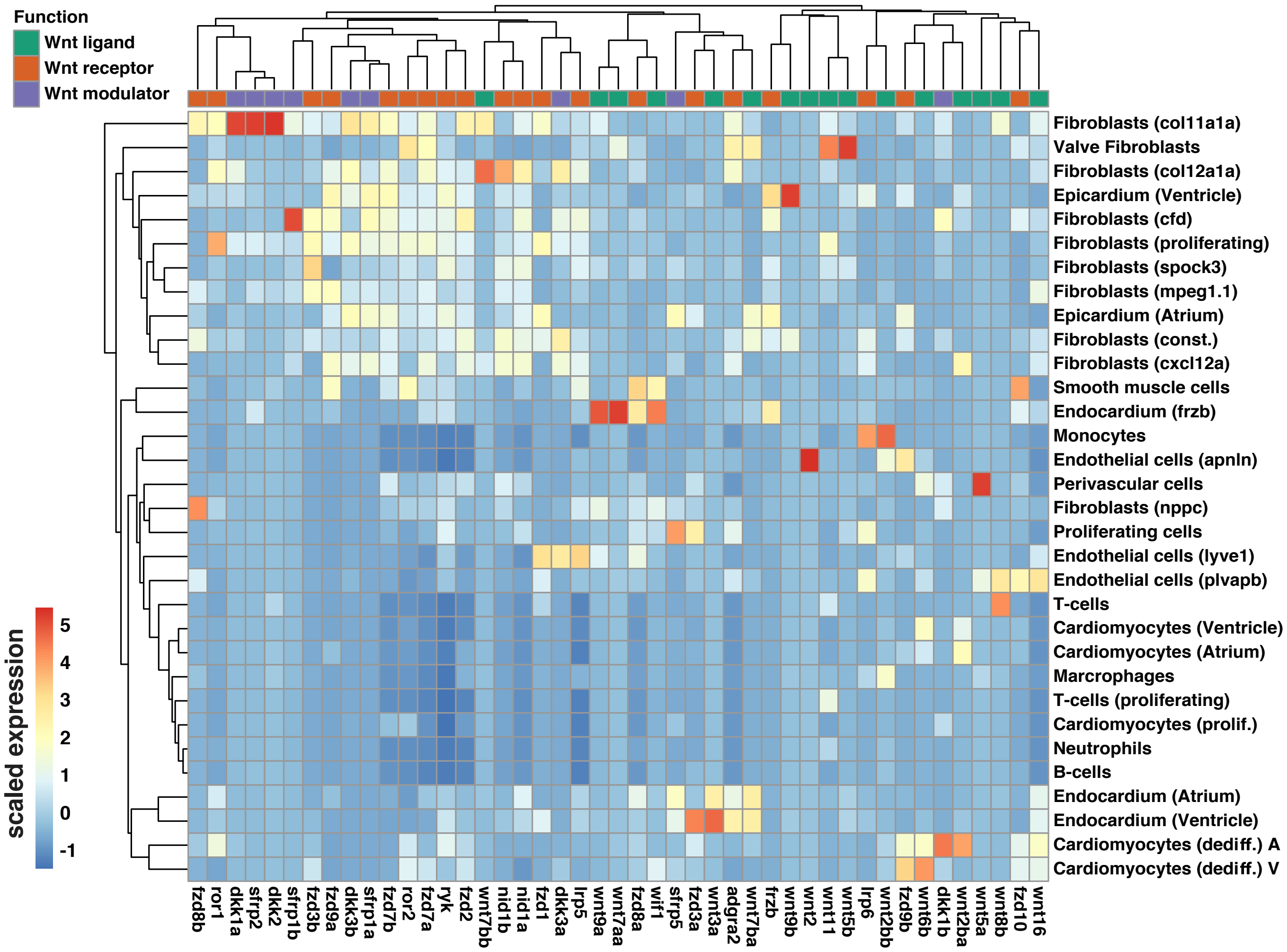


**Figure S19. Expression of Wnt signalling factors in the different cell types of the zebrafish heart.** Same analysis as in Fig. 5A, but with sub-cell-type resolution.

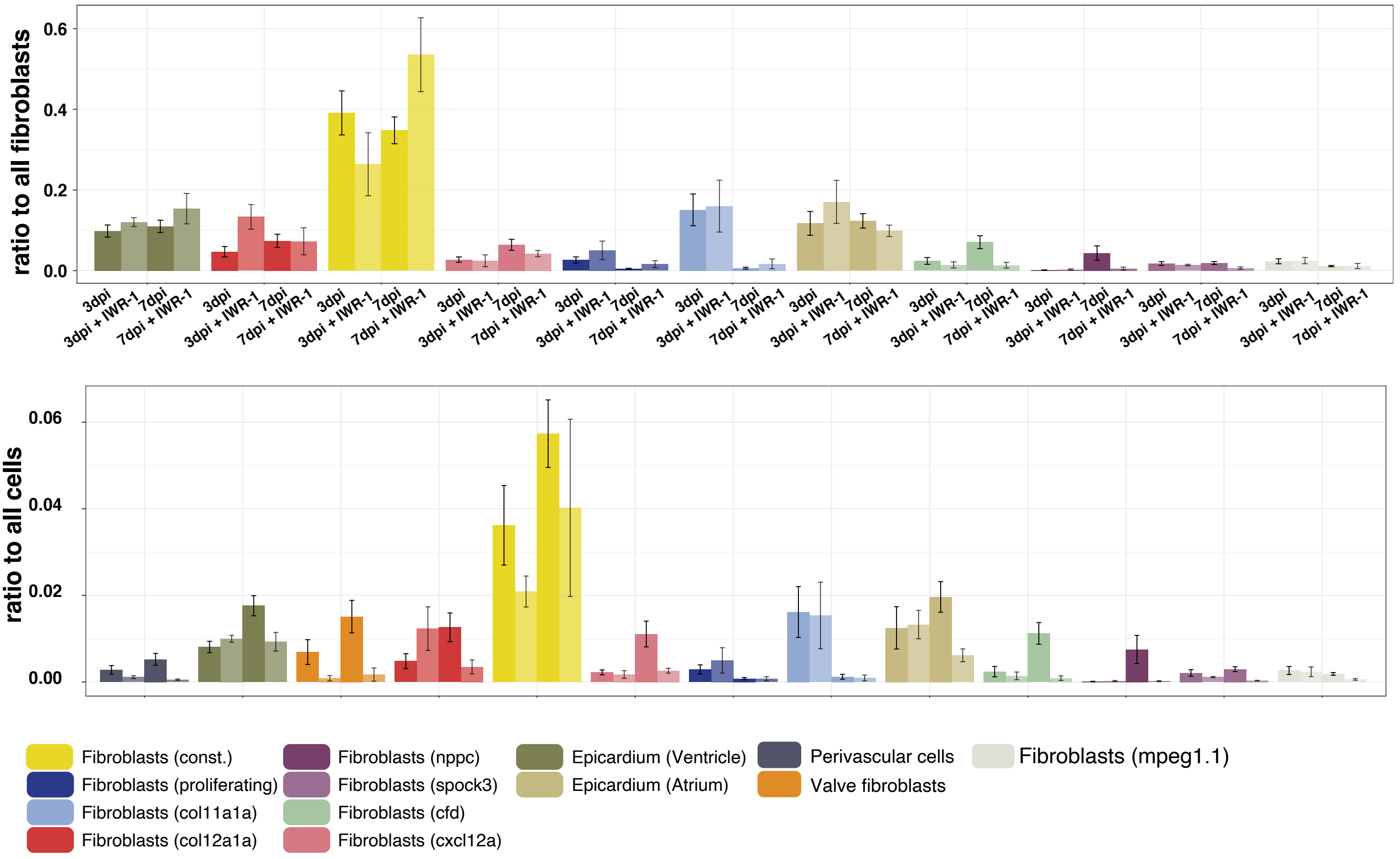


**Figure S20. Comparison of cell type abundance for all fibroblast subtypes at 3 and 7 dpi with or without Wnt inhibition.** Two types of normalization are shown: either cell numbers were normalized to total number of fibroblasts, or to total number of cells. Mean value across all replicates of the same treatment and timepoint is shown, error bars indicate standard error of the mean.
